# A plant virus, TYLCV, up-regulates an endocrine oxylipin signal in its insect vector, *Bemisia tabaci*, via the viral C2 virulence factor for viral transmission

**DOI:** 10.64898/2026.02.09.704802

**Authors:** Niayesh Shahmohammadi, Tae-Geun Song, Man-Cheol Son, Jiho Jeon, Donghee Lee, Eui-Joon Kil, Yonggyun Kim

## Abstract

To survive and efficiently transit between plant and insect hosts, circulative plant viruses have evolved sophisticated strategies to exploit insect vector factors. Tomato yellow leaf curl virus (TYLCV) is transmitted by *Bemisia tabaci* through a circulative and replicative pathway. In insects, C20 oxylipin (eicosanoid) and C18 oxylipin (EpOME) antagonistically regulate antiviral responses. Upon TYLCV infection, the intestinal apoptosis of *B. tabaci* facilitated the viral multiplication. The apoptosis was suppressed by eicosanoid but induced by EpOME. EpOME treatment also upregulated other proviral factors, including defensin, PGRP, and cathepsins, while eicosanoid signaling exerted opposite effects. TYLCV infection suppressed eicosanoid biosynthetic enzymes but induced a cytochrome P450 gene involved in EpOME biosynthesis, consistent with elevated EpOME levels in the viruliferous *B. tabaci* detected by LC-MS/MS. Individual RNA interference treatments specific to each of the TYLCV genes in the viruliferous insects revealed that only silencing of the viral C2 gene abolished EpOME-mediated proviral effects. These findings uncover a lipid-mediated mechanism by which TYLCV enhances vector competence to promote transmission.

**IMPORTANCE:** Various plant viruses depend on insect vectors for their horizontal transfer. Some of them exhibit a circulative and propagative transmission by multiplying the viral titers within the insects using the host machinery. Here is a fascinating manipulation of the host immunity by a plant virus, tomato yellow leaf curl virus (TYLCV), which uses the insect endocrine signals associated with immunity of its vector, *Bemisia tabaci*. Two types of oxylipins, eicosanoid and EpOME, antagonistically act to insect immunity, in which eicosanoid induces immune responses while EpOME, as an insect immune resolvin, suppresses them. TYLCV uses its C2 gene component as a virulent factor to induce the EpOME biosynthesis of *B. tabaci*. The elevated EpOME levels in the vector insect led to proviral responses by inducing intestinal apoptosis and selectively suppressing the immune-associated genes. These findings demonstrate the viral manipulation of the host endocrine signal for inducing proviral responses.

---

The silverleaf whitefly, *Bemisia tabaci* (Gennadius), is an insect pest damaging > 600 plant species by directly feeding and indirectly transmitting plant viruses classified in the genus *Begomovirus* (1, 2). It is highly invasive and now distributed across most subtropical and temperate regions worldwide, with a cryptic species complex consisting of at least 43 genetically distinct lineages, which are morphologically indistinguishable but can be discriminated by genetic and behavioral traits (3, 4). Among them, two invasive species, Middle East–Asia Minor 1 (MEAM1 = B biotype) and Mediterranean (MED = Q biotype), are particularly notorious for their ability to transmit plant viruses along with development of insecticide resistance (5, 6).

One of the most important viruses transmitted by *B. tabaci* is tomato yellow leaf curl virus (TYLCV, *Begomovirus coheni*), a member of *Begomovirus* with a single-stranded DNA genome (7). Phylogenetic studies have identified at least six TYLCV-related viruses, including TYLCV, tomato yellow leaf curl Sardinia virus (TYLCSV), tomato yellow leaf curl Axiarqua virus (TYLCAxV), tomato yellow leaf curl Malaga virus (TYLCMalV), tomato yellow leaf curl Mali virus (TYLCMV), and tomato leaf curl Sudan virus (ToLCSDV) (8). Among these, TYLCV is the most widespread and emergent species. TYLCV-induced disease is regarded as one of the most devastating plant diseases in temperate regions (9). Typical symptoms include yellowing and upward curling of leaflet margins, plant stunting, and flower abortion (10). Infected plants are less vigorous, yield fruits of reduced market value, and when infected during early growth stages, may experience complete crop loss. The first TYLCV genome was sequenced from Israel (TYLCV-IL) in 1991 (11), followed by the identification of a mild strain, TYLCV-Mld (12). Based on sequence identity, five major strains have been described, including Gezira (TYLCV-Gez), Iran (TYLCV-IR), Oman (TYLCV-OM), TYLCV-IL, and TYLCV-Mld. Over the past decades, whiteflies and begomoviruses together have emerged as major threats to vegetable production worldwide (13).

In response to viral infection, insects mount antiviral immune defenses essential for their survival. Insect immunity is innate and relies on pathogen recognition mechanisms that are genetically pre-programmed (14). Upon recognition, oxylipins act as key mediators of nonself signaling to modulate immune responses of two effector tissues, such as hemocytes and fat body, which perform cellular and humoral immune responses (15). Oxylipins are oxygenated derivatives of polyunsaturated fatty acids (PUFAs) and are broadly categorized into two subgroups in insects (16). Eicosanoids, including prostaglandins (PGs), leukotrienes, and epoxyeicosatrienoic acids, are C20 PUFAs that promote immune responses. Their biosynthesis is typically induced upon pathogenic infection through the activation of phospholipase A_2_ (PLA_2_), which liberates arachidonic acid (20:4n-6) and other C20 PUFAs from membrane phospholipids. In contrast, epoxyoctadecamonoenoic acids (EpOMEs), including vernolic acid (12,13-EpOME) and coronaric acid (9,10-EpOME), are C18 oxylipins that act as immune suppressors (17). They are synthesized by cytochrome P450 monooxygenases (CYPs) and degraded by soluble epoxide hydrolase (sEH) in different insect species (18, 19).

Pathogens often exploit oxylipin pathways to subvert host immunity. For example, the symbiotic bacteria *Xenorhabdus* and *Photorhabdus-*associated with entomopathogenic nematodes *Steinernema* and *Heterorhabditis*, respectively, produce secondary metabolites such as benzylideneacetone and non-ribosomal peptides that inhibit PLA_2_ activity, thereby blocking eicosanoid biosynthesis (20, 21). This suppression prevents the insect host from mounting immune responses, creating an immunocompromised environment that facilitates bacterial proliferation and nematode development within the insect cadaver (22). Similarly, tomato spotted wilt virus (TSWV) manipulates EpOME metabolism in its thrips vector, *Frankliniella occidentalis*, to antagonize eicosanoid-mediated antiviral responses (23, 24). Viral infection induces prostaglandin E_2_ (PGE_2_) biosynthesis, which mediates apoptosis in the midgut epithelium of thrips larvae and restricts viral replication (25). However, EpOMEs counteract this antiviral apoptotic response. A viral factor encoded by the NSs gene of TSWV has been shown to upregulate EpOME biosynthesis, thereby promoting viral replication (24). The proviral role of EpOMEs has also been demonstrated in entomopathogenic baculoviruses (26). These findings suggest that EpOME biosynthesis may represent a common molecular target exploited by different viral pathogens to suppress insect immunity.

This study was designed to test a hypothesis that TYLCV manipulates EpOME biosynthesis in its insect vector *B. tabaci* to facilitate circulative viral transmission. Specifically, we aimed to identify the biosynthetic machinery underlying the production of the two oxylipins and to evaluate their respective roles in TYLCV transmission.

## RESULTS

### Fatty acid metabolism is altered in whiteflies infected with TYLCV

TYLCV infected the intestine of *B. tabaci* after being fed (Fig. 1A). The viral infection was detected as early as 6 h in the midgut and hindgut, but not in the foregut visible in a thin thread-like structure (Fig. S1). In a time-course analysis, viral abundance increased in both the midgut and hindgut, as reflected by FISH signal intensity. Transcriptomic analysis (Table S3) of infected adults revealed that TYLCV infection altered the gene expression associated with major nutrients and their energy metabolism (Fig. 1B). KEGG pathway analysis on the transcriptome of *B. tabaci* adults after a 24-h AAP showed that one of the most noticeable changes after the viral infection was observed in fatty acid metabolism, in which its synthesis was active in nonviruliferous but not in viruliferous insects, while its modification was active in viruliferous but not in nonviruliferous insects, suggesting a substantial reprogramming in fatty acid metabolism associated with TYLCV infection.

**Fig 1.**
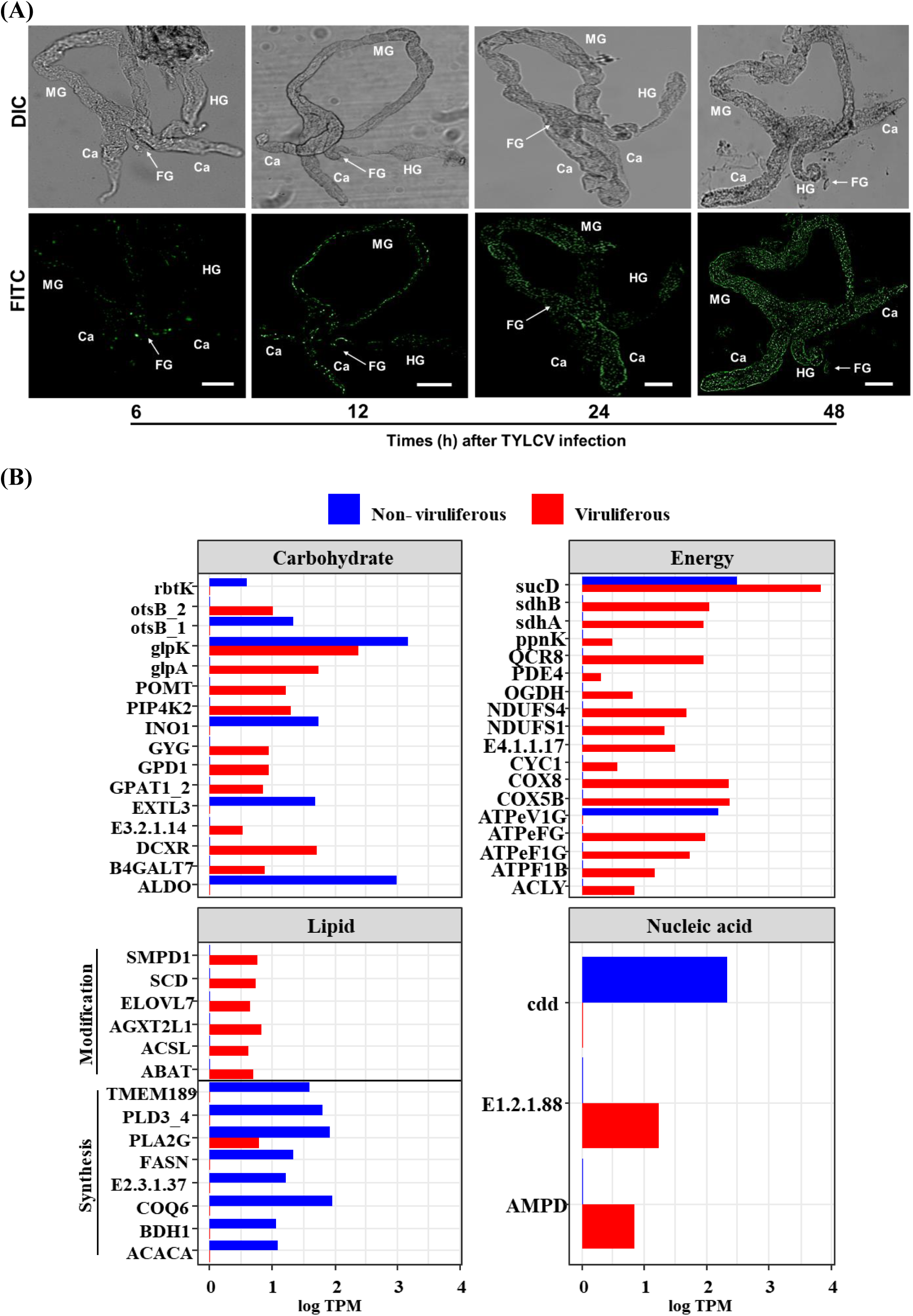
TYLCV multiplication in the intestine of *B. tabaci* and alteration of host metabolism. (A) Progressive increase in TYLCV accumulation in the intestine of *B. tabaci* during infection. The viral localization was monitored by FISH analysis in the intestine of adult whiteflies at 6, 12, 24, and 48 h post-infection: ‘Ca’ for caeca, ‘FG’ for foregut, ‘MG’ for midgut, and ‘HG’ for hindgut. The intestine was visualized using differential interference contrast (DIC), and TYLCV signals were detected using fluorescein isothiocyanate (FITC) under a fluorescence microscope (DM2500; Leica, Wetzlar, Germany). Scale bar = 0.1 mm. (B) Comparative expression profiles of genes involved in different metabolic pathways using KEGG analysis between viruliferous and non-viruliferous whiteflies. Expression levels are presented as log-transformed transcripts per million (log TPM). The full gene names of the abbreviated accessions are listed in Table S4.

### Two oxylipins oppositely regulate TYLCV titers in whiteflies

Two types of oxylipins, PGE_2_ and EpOMEs, are derived from phospholipids through oxygenation pathways (Fig. 2A). The effects of these oxylipins on TYLCV titers were examined in whiteflies. The precursors, arachidonic acid (AA) and linoleic acid (LA), exhibited opposite effects on viral accumulation, with AA significantly reducing and LA increasing viral titers (Fig. 2B). Among eicosanoids resulting from AA, three PGs were compared for their anti-viral activities against TYLCV because they were detected in other insects (16), PGE_2_ was the most potent to in suppressing the viral titers, whereas PGF_2α_ had no detectable effect (Fig. 2C). In contrast, both EpOME isomers significantly enhanced TYLCV titers, with 12,13-EpOME showing a stronger effect than 9,10-EpOME (Fig. 2D).

**Fig 2.**
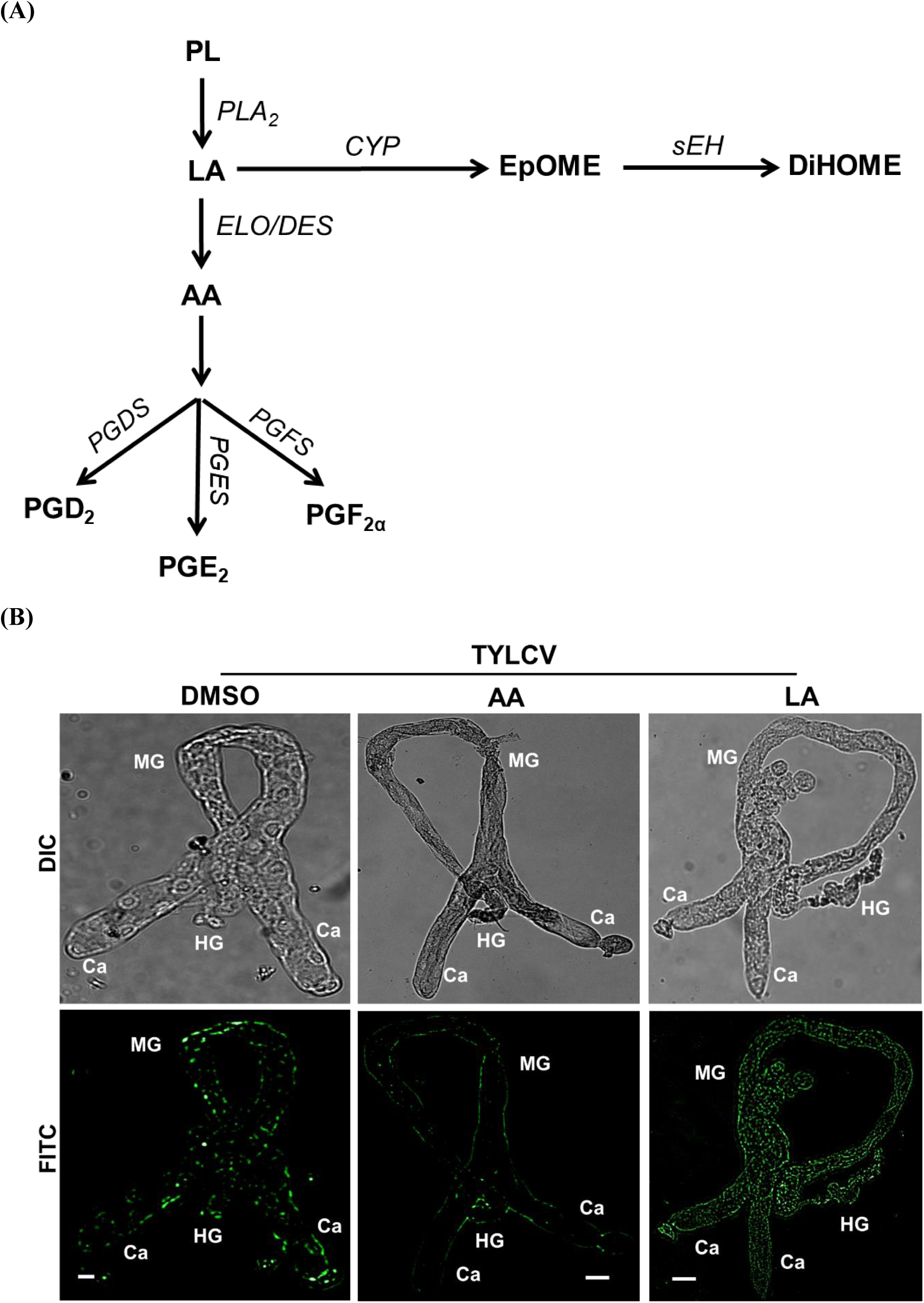
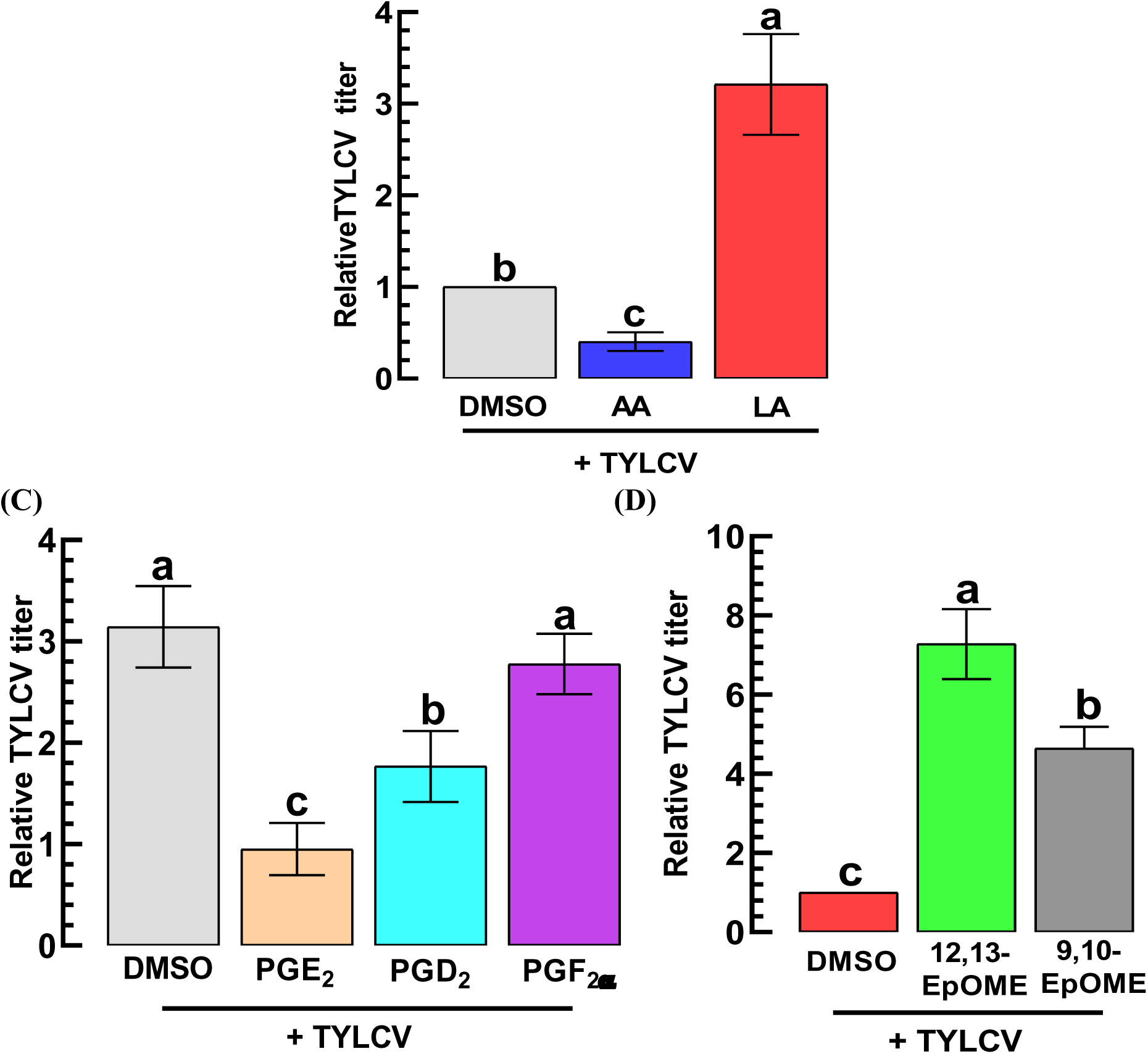
Opposite roles of two oxylipins (PGE_2_ and EpOME) in manipulating TYLCV multiplication. (A) A diagram indicating the biosynthetic pathways for the oxylipins. Phospholipid (‘PL’) is catalyzed by phospholipase A_2_ (‘PLA_2_’) to release linoleic acid (‘LA’). LA is then oxygenated by cytochrome P450 monooxygenase (‘CYP’) to produce EpOME, which is degraded into DiHOME by soluble epoxide hydroxylase (‘sEH’). LA is also elongated and desaturated by elongase (‘ELO’) and desaturase (‘DES’), respectively, to produce arachidonic acid (‘AA’). AA is then further oxygenated to different prostaglandins (PGs) by PGD_2_ synthase (‘PGDS’), PGE_2_ synthase (‘PGES’), and PGF_2α_ synthase (‘PGFS’). (B) Effects of oxylipin precursors (AA and LA) on TYLCV multiplication in the intestine assessed by FISH. The intestine was visualized using differential interference contrast (‘DIC’), and TYLCV signals were detected using fluorescein isothiocyanate (‘FITC’) under a fluorescence microscope (DM2500; Leica, Wetzlar, Germany): ‘Ca’ for caeca, ‘MG’ for midgut, and ‘HG’ for hindgut. Scale bar = 0.1 mm. The viral titers were assessed in the whole-body using qPCR at 24 h after treatment. (C) Effects of three different PGs (0.1 μg/mL) on TYLCV multiplication assessed by qPCR at 24 h after treatment. (D) Effects of two EpOMEs (1 μg/mL) on TYLCV multiplication assessed by qPCR at 24 h after treatment. All compounds were administered to adult whiteflies by a membrane feeding together along with TYLCV. ‘DMSO’ represents a solvent control. At 24 h after treatment, TYLCV titers were quantified by qPCR. Each treatment was performed with three biological replicates. Different letters above the standard deviation bars indicate significant differences among treatments at Type I error = 0.05 (LSD test).

### Up-regulation of PGE_2_ signaling components in TYLCV-infected whiteflies

PGE_2_ biosynthesis is initiated by the catalytic activity of PLA_2_, which releases AA from membrane phospholipids (Fig. 3A). Pharmacological inhibition of PLA_2_ using dexamethasone (DEX) resulted in a significant increase in TYLCV titers in the whiteflies. Similarly, inhibition of cyclooxygenase (COX) or lipoxygenase (LOX) also increased the viral titers, with naproxen (NAP) treatment specific to COX being more potent than esculetin (ESC) treatment specific to LOX.

**Fig 3.**
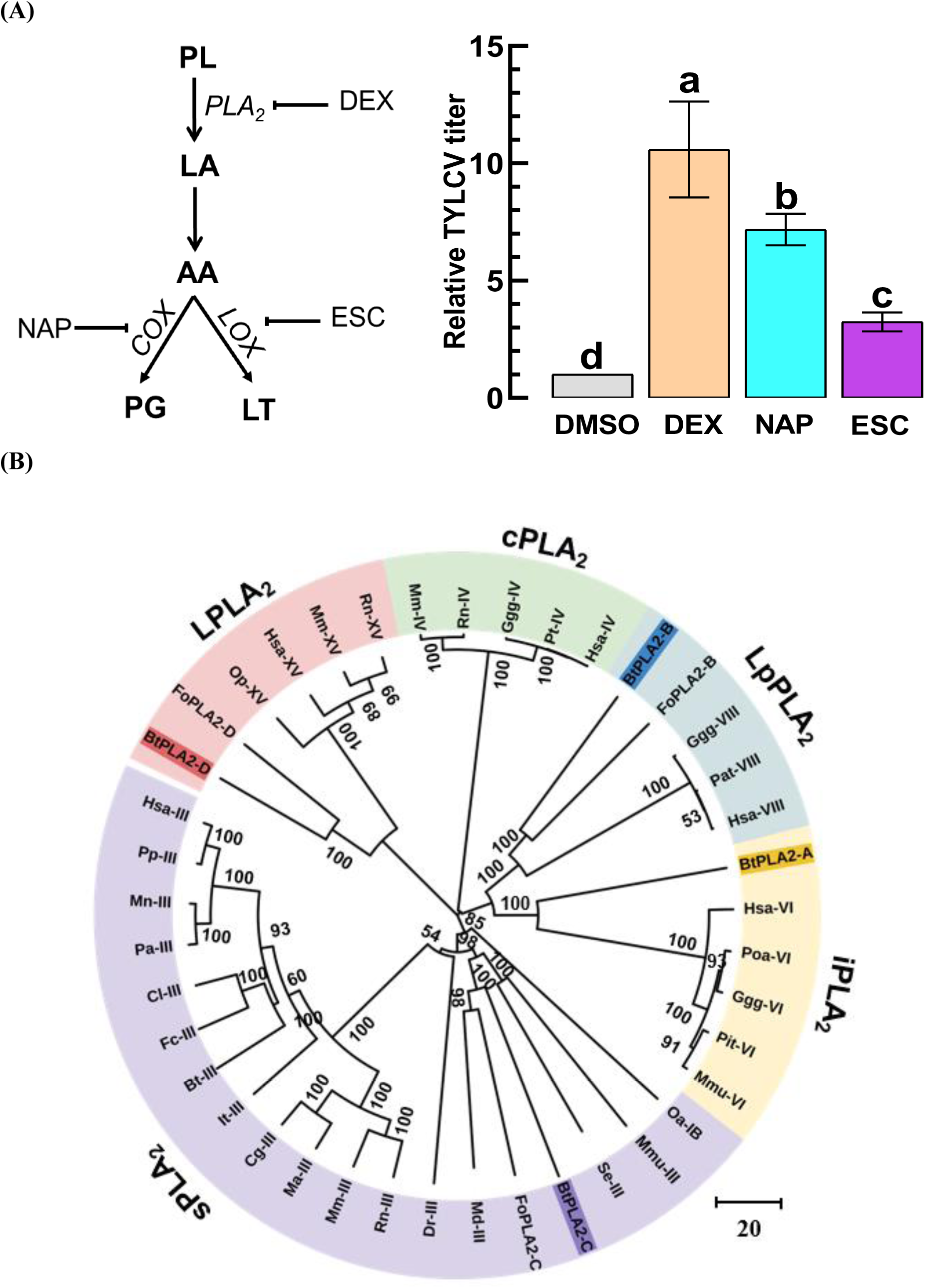
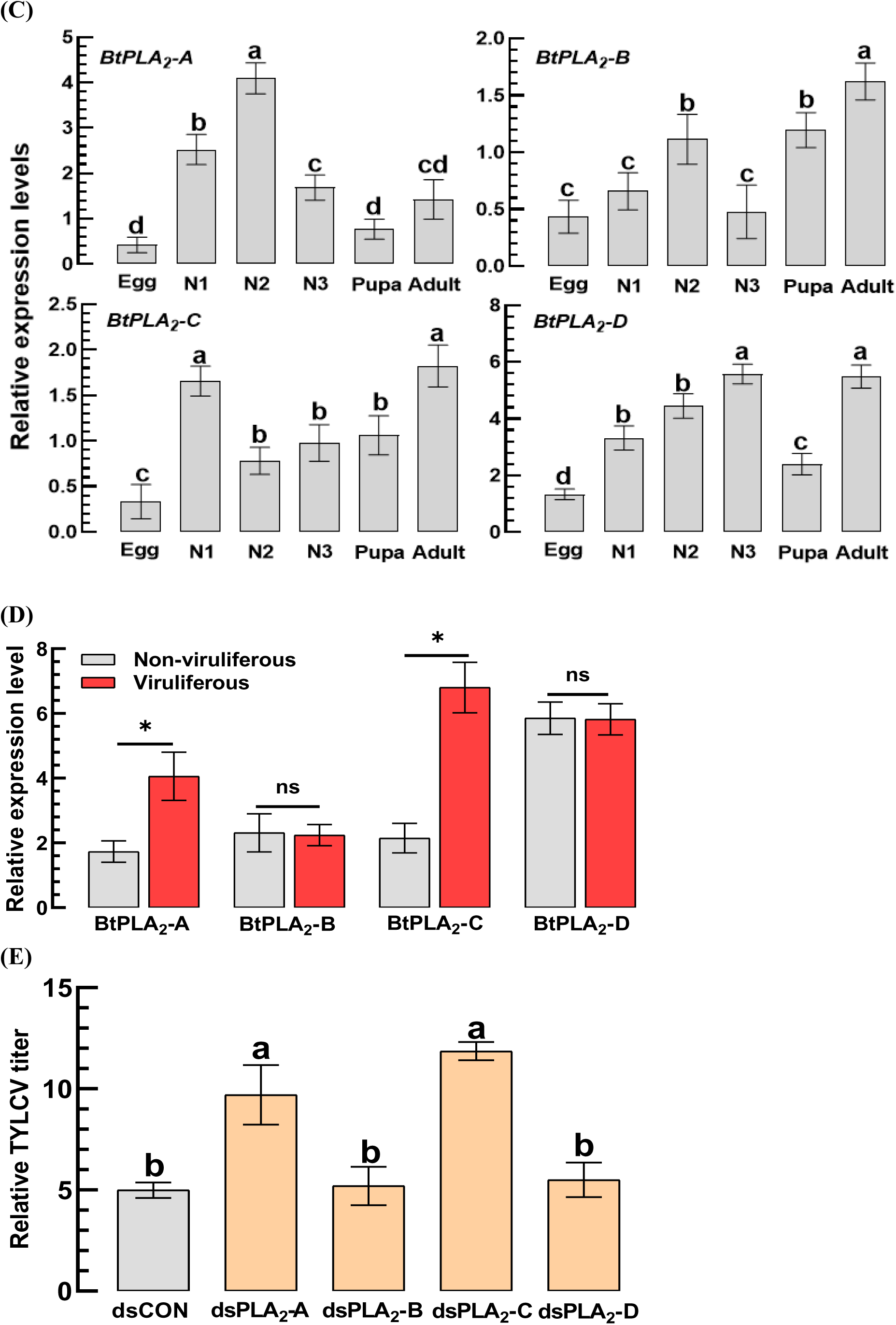
Phospholipase A_2_ (PLA_2_) genes of *B. tabaci* and their antiviral activity against TYLCV. (A) Influence of three inhibitors (1 μg/mL) specific to eicosanoid biosynthesis on TYLCV multiplication in *B. tabaci*: dexamethasone (‘DEX’) specific to PLA_2_, naproxen (‘NAP’) specific to cyclooxygenase (‘COX’), and esculetin (‘ESC’) specific to lipoxygenase. ‘DMSO’ represents solvent control. (B) A phylogenetic analysis of four PLA_2_ genes (*BtPLA_2_-A* ∼ *BtPLA_2_-D*) encoded in the *B. tabaci* genome, along with other PLA_2_ genes separately identified into 15 Groups (I-XV), which are also classified into five different PLA_2_ types: secretory (sPLA_2_), lysosomal (LPLA_2_), lipoprotein-associated (LpPLA_2_), Ca^2+^-independent cellular (iPLA_2_), and Ca^2+^-dependent cellular (cPLA_2_). (C) Developmental expression profiles of the four PLA_2_ genes from egg to adult. ‘N1-N3’ represents three nymphal instars. (D) Influence of TYLCV infection on the expression levels of the four PLA_2_ genes. qPCR was analyzed at 24 h after viral infection in adults. (E) Effects of four individual RNA interference (RNAi) treatments using gene-specific dsRNAs (dsPLA_2_-A ∼ dsPLA_2_-D, 500 µg/mL) against the four PLA_2_ genes on TYLCV titers of the viruliferous adults. ‘dsCON’ indicates a control dsRNA. After 24 h, the viral titers were quantified using qPCR. Different letters or asterisks above the standard deviation bars indicate significant differences among treatments at Type I error = 0.05 (LSD test). ‘ns’ represents no significant difference.

Four *PLA_2_* genes were predicted from the *B. tabaci* genome and classified through a phylogenetic analysis into distinct PLA_2_ groups: *BtPLA_2_-A* (*iPLA_2_*, Group VI), *BtPLA_2_-B* (*LpPLA_2_*, Group VIII), *BtPLA_2_-C* (*sPLA_2_*, Group III), and *BtPLA_2_-D* (*LPLA_2_*, Group XV) (Fig. 3B). All four genes were expressed across developmental stages from egg to adult (Fig. 3C). Upon TYLCV infection, *BtPLA_2_-A* and *BtPLA_2_-C* were significantly up-regulated, whereas the other two genes showed no significant changes (Fig. 3D). RNAi-mediated silencing of *BtPLA_2_-A* or *BtPLA_2_-C* increased the viral titers, while knockdown treatments of the other two *PLA_2_*genes did not change the viral titers (Fig. 3E).

PGE_2_ synthase (*PGES2*) catalyzes the conversion of PGH_2_ to PGE_2_ following AA oxygenation by COX (Fig. 4A). PGE_2_ exerts its effects through binding to the PGE_2_ receptor (*PGE_2_R*). Both *Bt-PGES2* and *Bt-PGE_2_R* were predicted through orthologous phylogenetic analyses and clustered with hemipteran orthologs, distinct from other insect groups (Fig. 4B). These genes were expressed throughout all developmental stages (Fig. 4C) and significantly up-regulated following TYLCV infection (Fig. 4D). RNAi treatment specific to either *Bt-PGES2* or *Bt-PGE_2_R* significantly increased the viral titers in viruliferous whiteflies (Fig. 4E). Together, these results indicate that TYLCV infection induces the antiviral response of *B. tabaci* by elevating the eicosanoid biosynthesis, leading to PGE_2_ production that suppresses viral accumulation.

**Fig 4.**
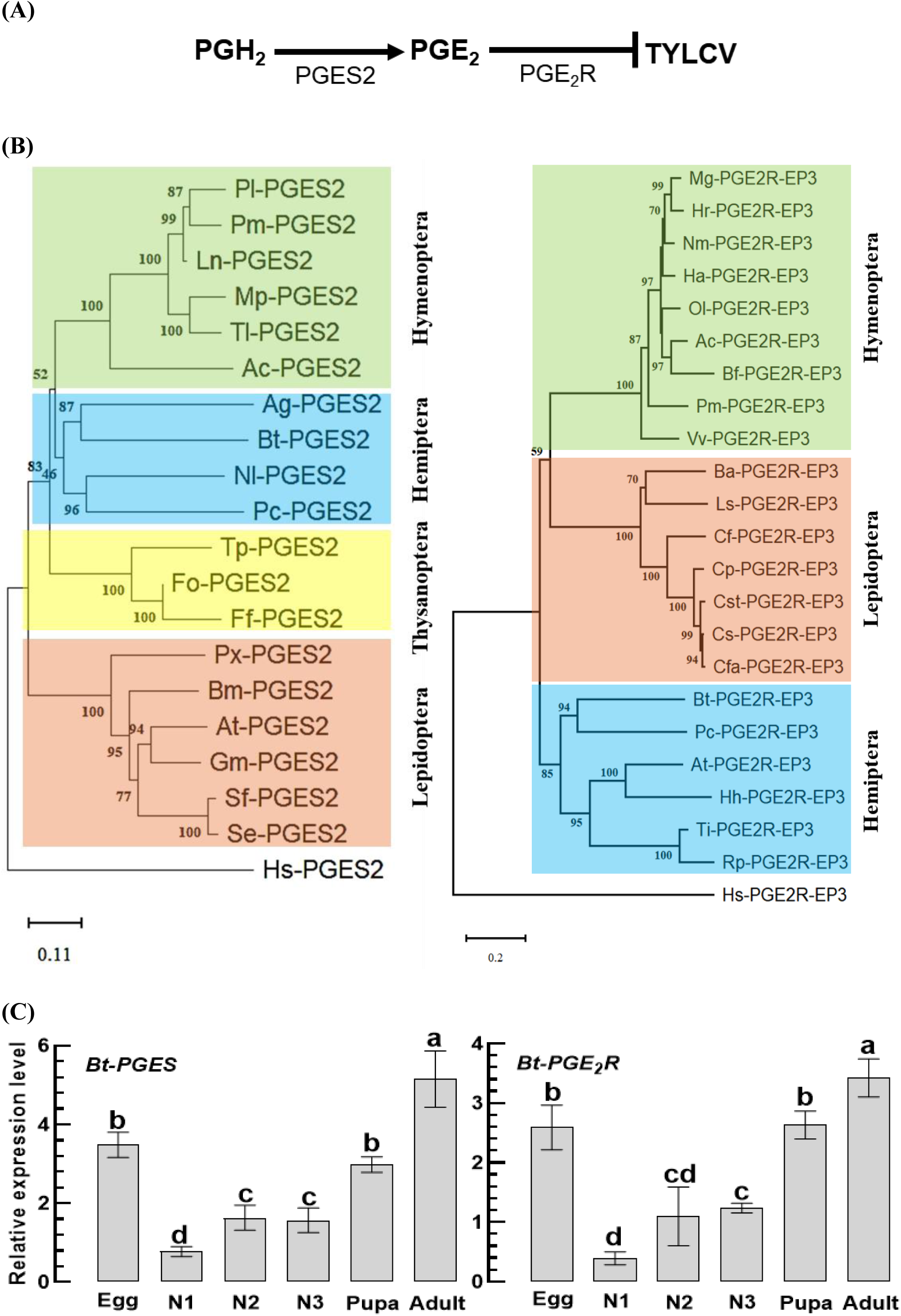
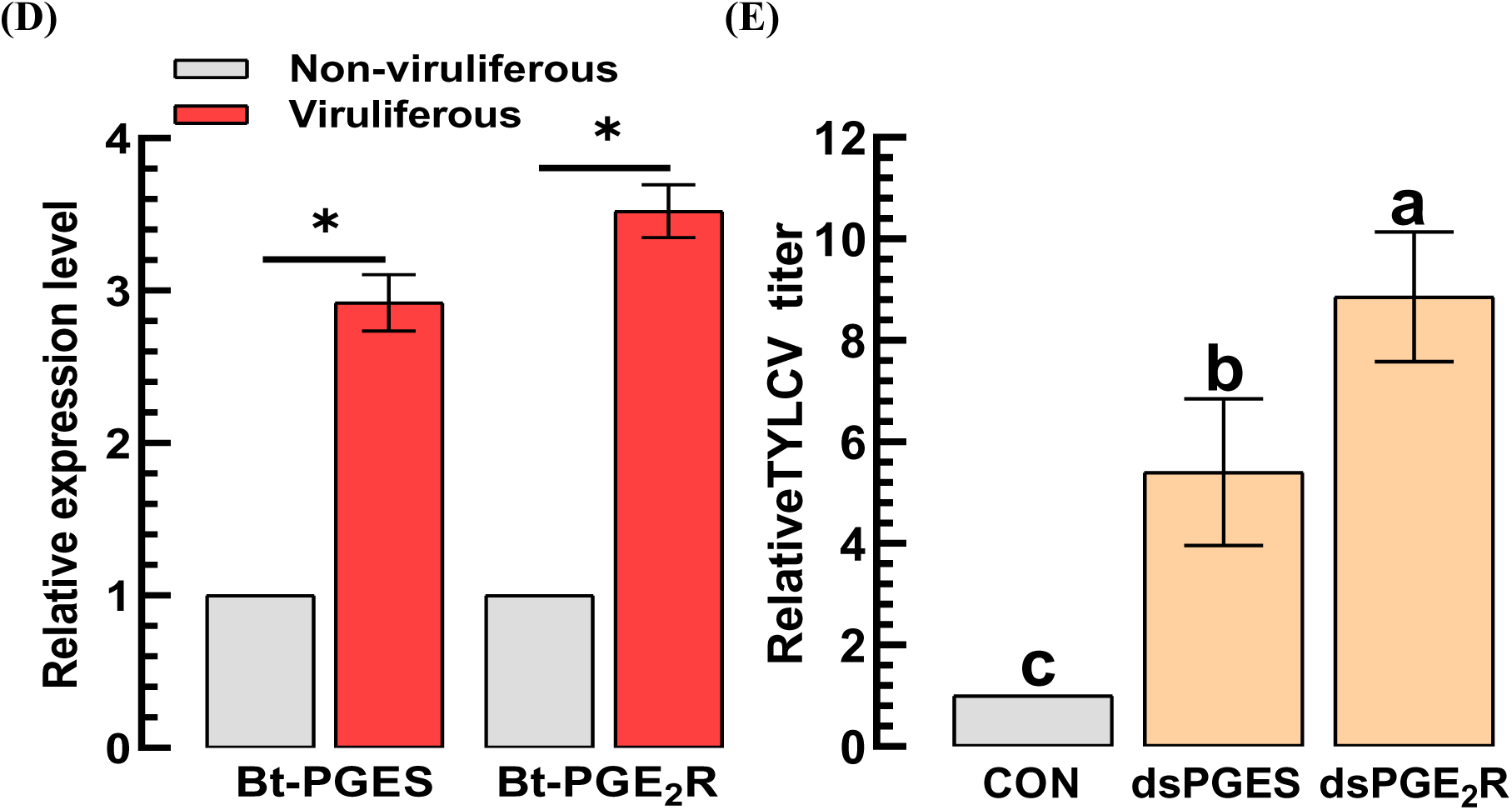
PGE_2_ synthase (Bt-PGES2)/receptor (Bt-PGE_2_R) of *B. tabaci* and their antiviral activities against TYLCV. (A) A diagram indicating an antiviral pathway of PGE_2_, in which PGE_2_ is synthesized from PGD_2_ by PGES2 and mediates an antiviral response via its PGE_2_R. (B) Phylogenetic analyses of *Bt-PGES* and *Bt-PGE_2_R* encoded in the *B. tabaci* genome along with their orthologous genes. (C) Developmental expression profiles of the two genes from egg to adult. ‘N1-N3’ represents three nymphal instars. (D) Up-regulation of the two genes in response to TYLCV infection. (E) Influence of two individual RNA interference (RNAi) treatments using gene-specific dsRNAs (dsPGES and dsPGE_2_R, 500 µg/mL) against *Bt-PGES* or *Bt-PGE_2_R* genes on TYLCV titers of the viruliferous adults. ‘dsCON’ indicates a control dsRNA. After 24 h, the viral titers were quantified using qPCR. Different letters or asterisks above the standard deviation bars indicate significant differences among treatments at Type I error = 0.05 (LSD test).

### Up-regulation of EpOME biosynthesis in TYLCV-infected whiteflies

Two EpOME isomers were detected in *B. tabaci* using LC-MS/MS analysis (Fig. 5A). Their concentrations increased more than two-fold following TYLCV infection. Inhibition of EpOME degradation to DiHOME using AUDA significantly increased viral titers in viruliferous whiteflies (Fig. 5B). In addition, supplementation with an EpOME analog (AS56, 19) significantly enhanced viral accumulation. These results support the biosynthetic and degrading pathways of EpOME in *B. tabaci*.

**Fig 5.**
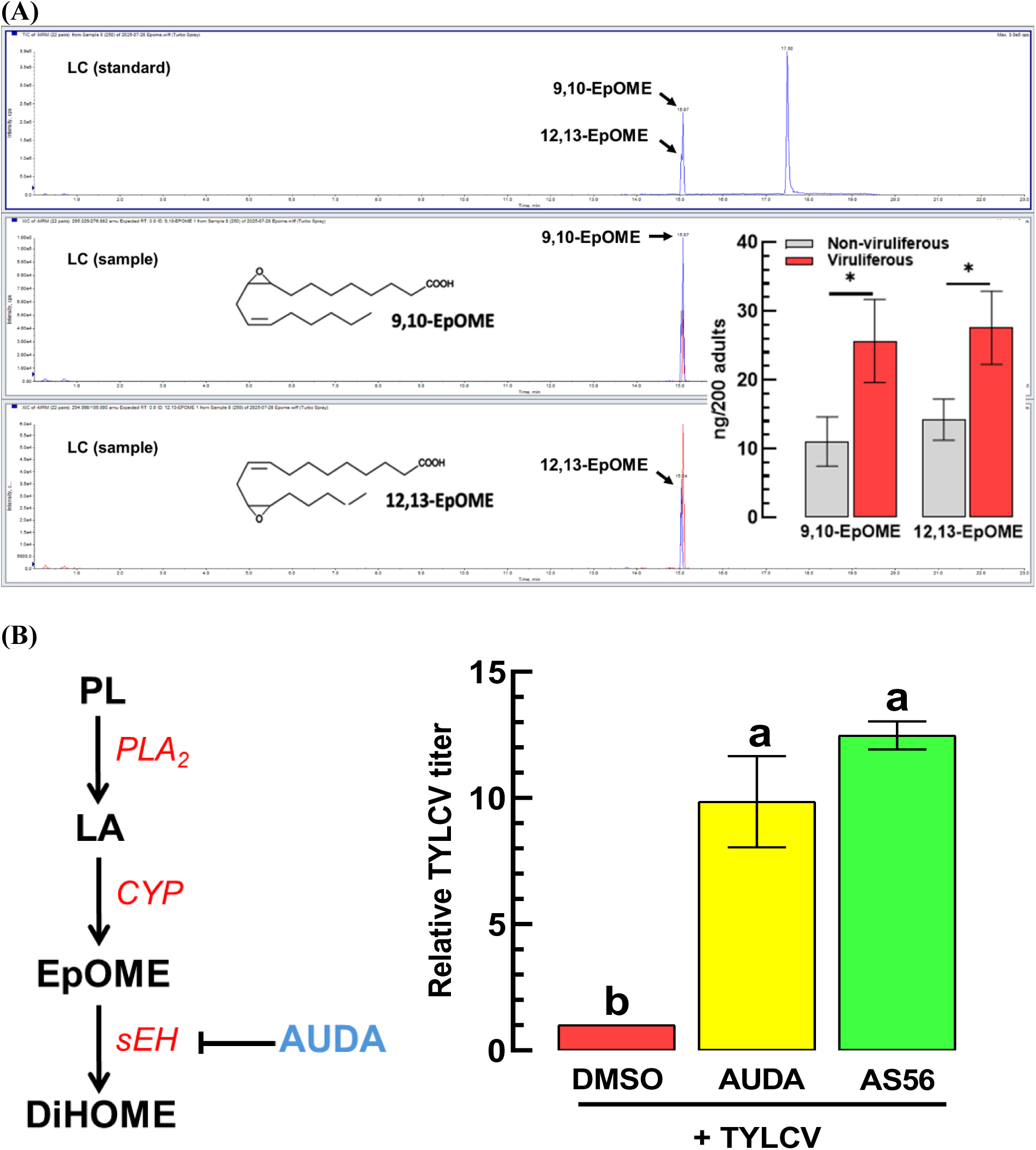
Up-regulation of EpOME levels in viruliferous *B. tabaci* and proviral activity for TYLCV. (A) Quantification of EpOMEs using LC–MS/MS. Two EpOME peaks are denoted in the chromatograms. (B) Proviral activity of EpOME for TYLCV. An inhibitor (‘AUDA’, 10 μg/mL) specific to soluble epoxide hydrolase (‘sEH’) or an EpOME analog (AS56, 0.1 μg/mL) was fed to the viruliferous *B. tabaci*. ‘DMSO’ represents a solvent control. After 24 h, the viral titers were quantified by qPCR. Different letters or asterisks above the standard deviation bars indicate significant differences among treatments at Type I error = 0.05 (LSD test).

To identify EpOME biosynthetic enzymes, cytochrome P450 (*CYP*) homologs known for their EpOME-biosynthetic function were phylogenetically analyzed (Fig. 6A). Three *CYP* genes (*Bt-CYP5*, *Bt-CYP6*, and *Bt-CYP7*) clustered with known EpOME synthases and were expressed across all developmental stages (Fig. 6B). Among them, only *Bt-CYP6* was significantly up-regulated following TYLCV infection (Fig. 6C). RNAi knockdown experiments demonstrated that silencing *Bt-CYP6* significantly reduced viral titers, whereas knockdown treatments of the other *CYP* genes had no effect (Fig. 6D), indicating a specific role for *Bt-CYP6* in EpOME biosynthesis during viral infection.

**Fig 6.**
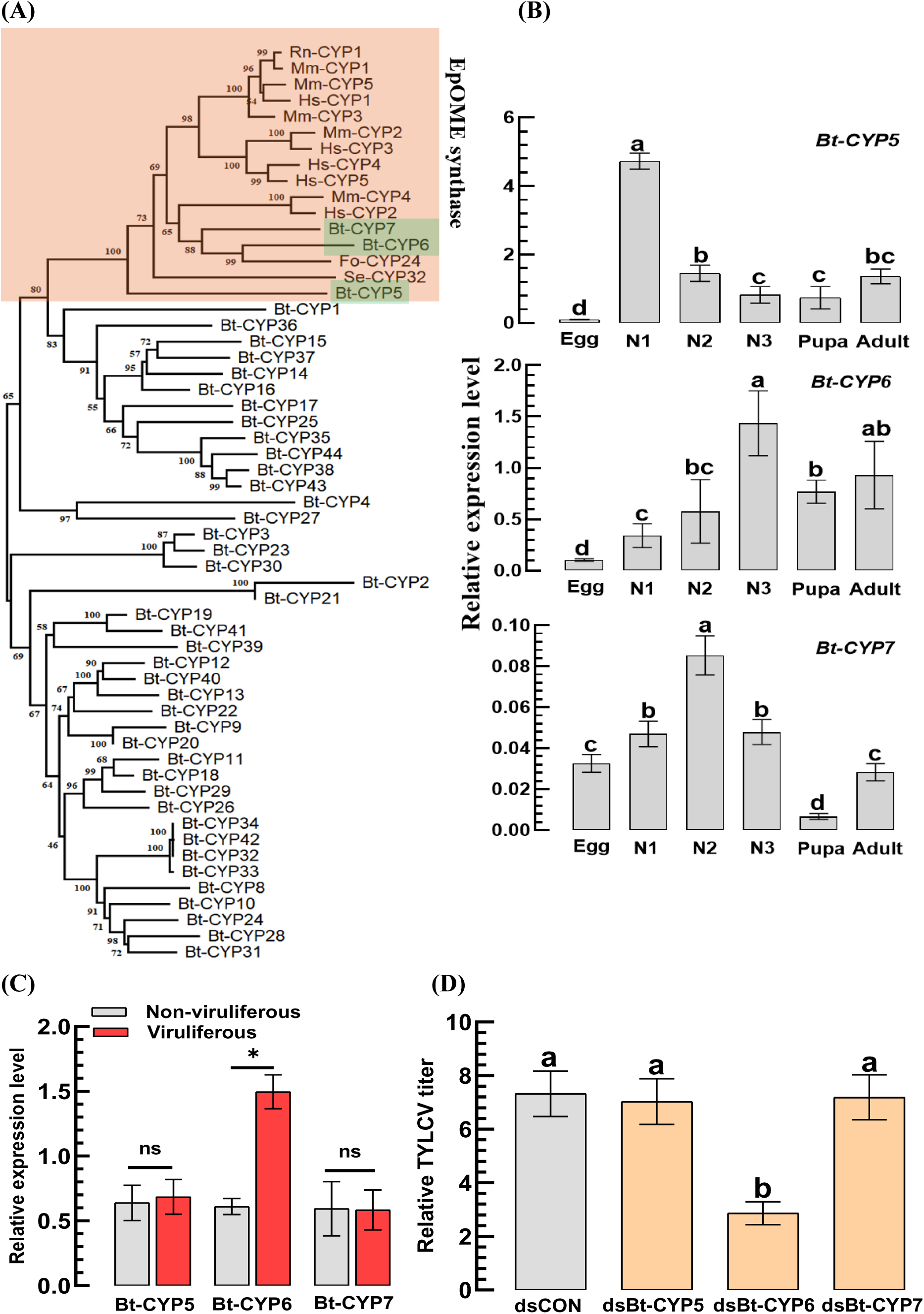
Cytochrome P450 monooxygenases (Bt-CYPs) of *B. tabaci* and their proviral activities for TYLCV. (A) Phylogenetic analysis of 44 *Bt-CYP* genes encoded in the *B. tabaci* genome, along with other EpOME synthases (*Fo-CYP24*, *Se-CYP32*, and *Hs-CYP2*). (B) Developmental expression profiles of the three CYP genes from egg to adult. ‘N1-N3’ represents three nymphal instars. (C) Influence of TYLCV infection on the expression levels of the three CYP genes. qPCR was analyzed at 24 h after viral infection in adults. (D) Influence of three individual RNA interference (RNAi) treatments using gene-specific dsRNAs (dsBt-CYP5 ∼ dsBt-CYP7, 500 µg/mL) against the three CYP genes on TYLCV titers of the viruliferous adults. ‘dsCON’ indicates a control dsRNA. After 24 h, the viral titers were quantified using qPCR. Different letters or asterisks above the standard deviation bars indicate significant differences among treatments at Type I error = 0.05 (LSD test).

EpOME-degrading enzymes were identified through a phylogenetic analysis, revealing four epoxide hydrolase genes: one membrane-bound juvenile hormone epoxide hydrolase (*JHEH*) and three soluble epoxide hydrolases (*sEH1*–*sEH3*) (Fig. 7A). All genes were expressed across developmental stages (Fig. 7B). Upon TYLCV infection, expression of *Bt-sEH1* ∼ *Bt-sEH3* was significantly suppressed, whereas *Bt-JHEH* expression remained unchanged (Fig. 7C). RNAi-mediated silencing of *Bt-sEH2* or *Bt-sEH3* significantly increased viral titers, while knockdown of *Bt-sEH1* or *Bt-JHEH* had no effect (Fig. 7D), indicating that *Bt-sEH2* and *Bt-sEH3* regulate EpOME levels during TYLCV infection.

**Fig 7.**
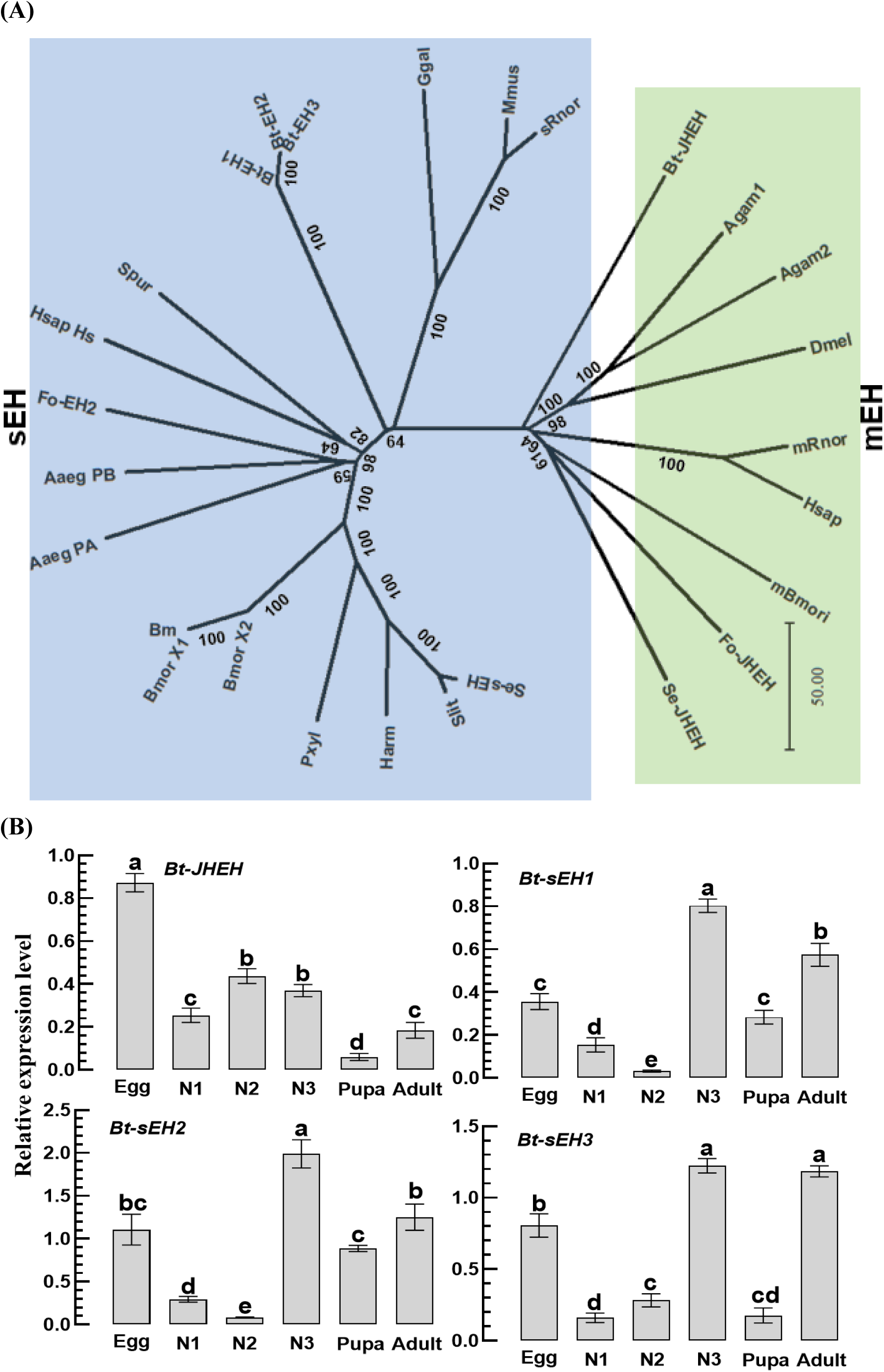
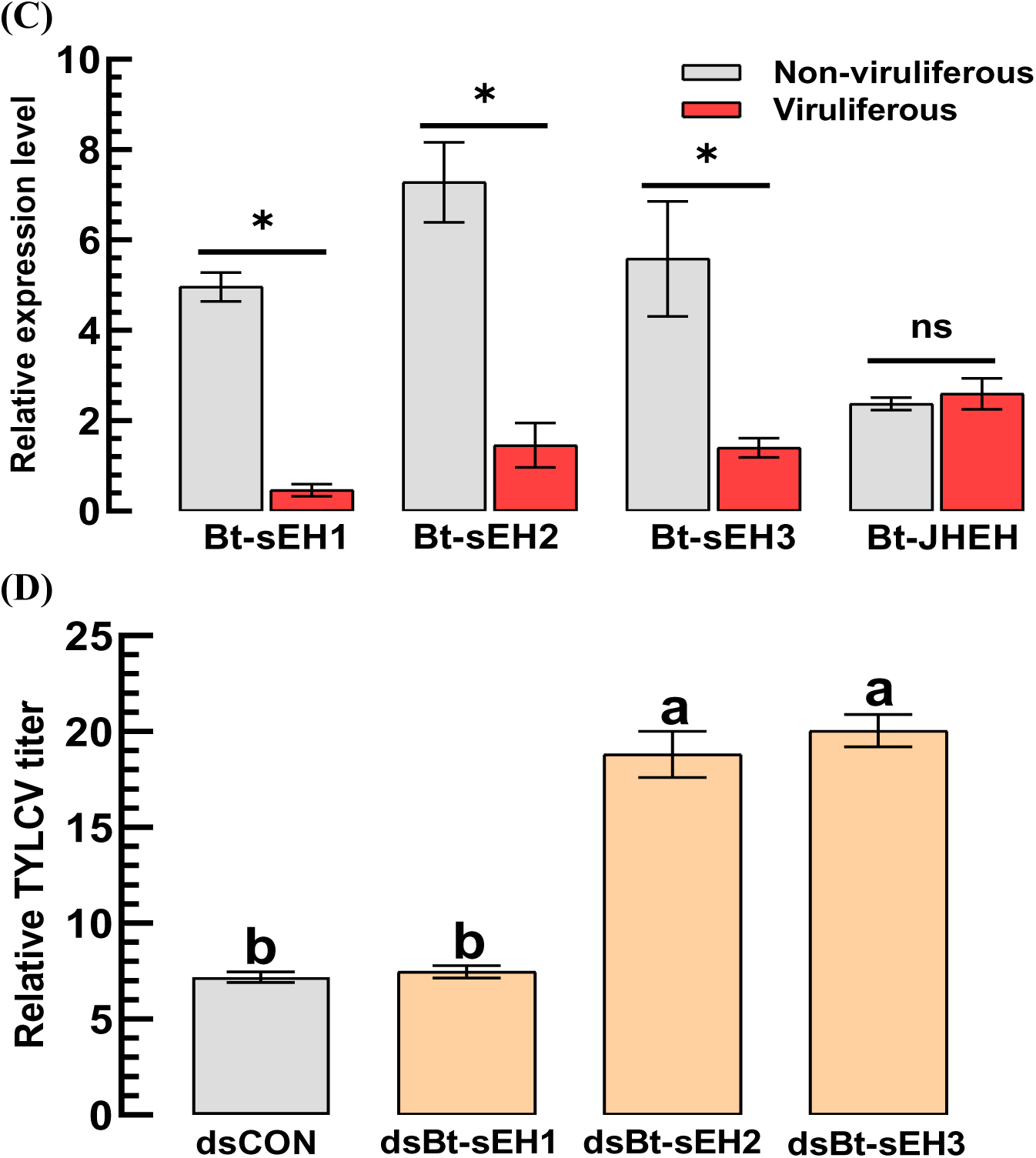
An EpOME-degrading enzyme, soluble epoxide hydrolase (Bt-sEH) of *B. tabaci* and its proviral activities for TYLCV. (A) Phylogenetic analysis of four epoxide hydrolase (*Bt-sEH1 ∼ sEH3, Bt-JHEH*) genes encoded in the *B. tabaci* genome along with other sEH and JHEH genes. (B) Developmental expression profiles of the four epoxide hydrolase genes from egg to adult. ‘N1-N3’ represents three nymphal instars. (C) Influence of TYLCV infection on the expression levels of the four epoxide hydrolase genes. qPCR was analyzed at 24 h after viral infection in adults. (D) Influence of three individual RNA interference (RNAi) treatments using gene-specific dsRNAs (ds-sEH1 ∼ ds-sEH3, 500 µg/mL) against the three sEH genes on TYLCV titers of the viruliferous adults. ‘dsCON’ indicates a control dsRNA. After 24 h, the viral titers were quantified using qPCR. Different letters or asterisks above the standard deviation bars indicate significant differences among treatments at Type I error = 0.05 (LSD test).

These results indicate the EpOME-associated metabolic genes of *B. tabaci*, in which *Bt-CYP6* is associated with EpOME biosynthesis and *Bt-sEH2*/*sEH3* are associated with EpOME degradation.

### Sequential expression profile of eicosanoid and EpOME biosynthesis-associated genes in *B. tabaci* after TYLCV infection

Two oxylipins (PGE_2_ and EpOME) exhibited antagonistic activities against TYLCV in *B. tabaci*. To examine their temporal production in *B. tabaci* after TYLCV infection, their synthetic genes were monitored in their expressions at different time points post-inoculation (Fig. 8). Three genes (*BtPLA_2_-A*, *BtPLA_2_-C*, and *Bt-PGES2*) associated with PGE_2_ biosynthesis were up-regulated immediately after the viral infection and showed their maximal expression at 1-day post-infection. In contrast, the EpOME biosynthesis gene (*Bt-CYP6*) exhibited relatively later expression patterns, with its maximal expression peaking at 2 ∼ 3 day post-infection. Interestingly, the EpOME-degradation genes (*Bt-sEH2*/*sEH3*) were significantly suppressed at 2 ∼ 3 days post-infection. These temporal patterns of gene expressions suggest differential regulation of PGE_2_ and EpOME biosynthesis during viral infection in *B. tabaci*.

**Fig 8.**
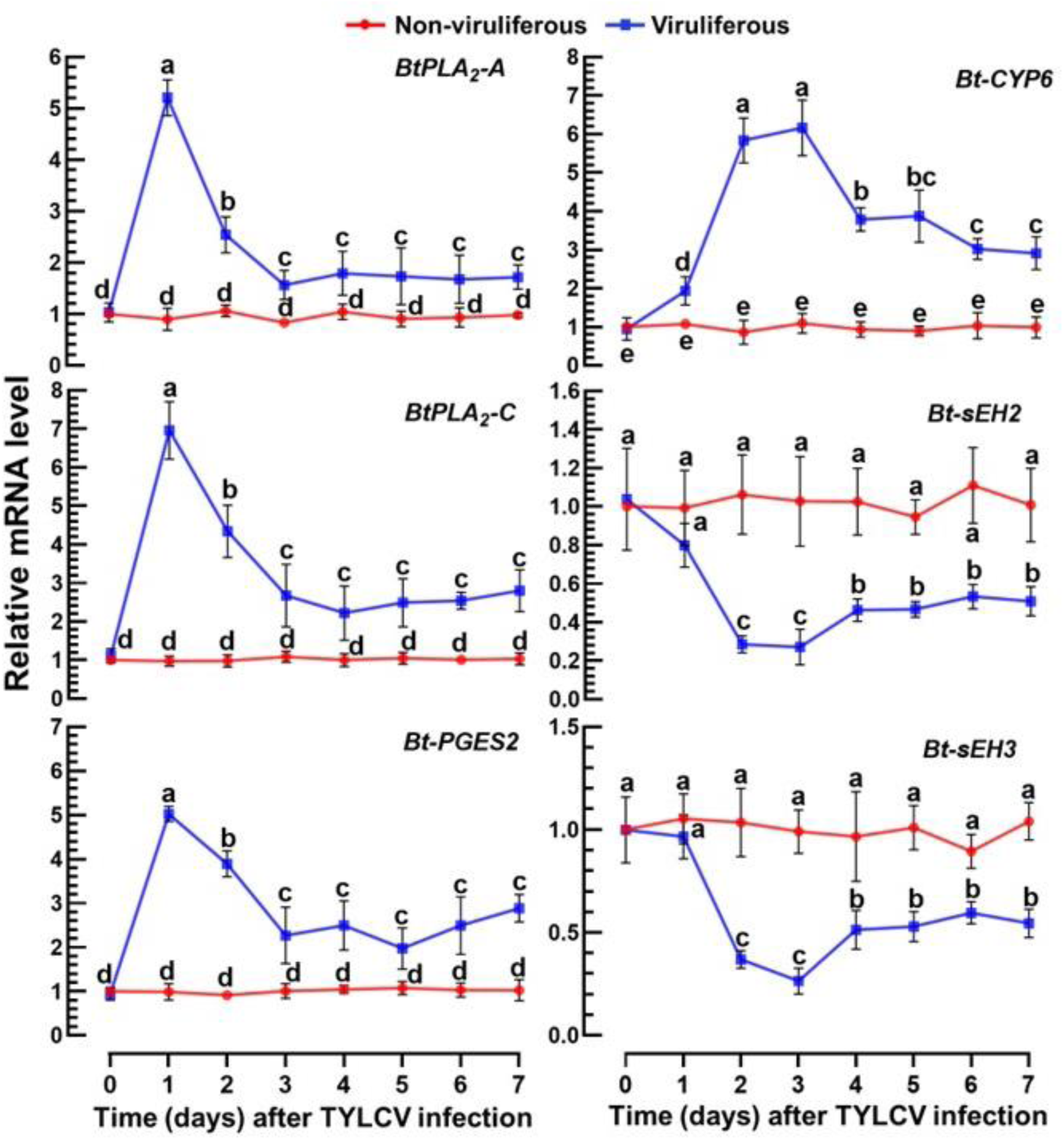
Temporal regulation of two oxylipin biosynthesis pathways in *B. tabaci* after TYLCV infection. Expression analyses were performed for seven days after viral infection of three genes (BtPLA_2_-A, BtPLA_2_-C, and Bt-PGES2) for PGE_2_ biosynthesis and three other genes (Bt-CYP6. Bt-sEH2, and Bt-sEH3) for EpOME biosynthesis. Different letters above the standard deviation bars indicate significant differences among treatments in each gene at Type I error = 0.05 (LSD test).

### Validation of EpOME as a proviral signal using two TYLCV strains

To support the role of EpOME as a proviral factor for TYLCV, two viral strains with different virulence were tested for their ability to induce EpOME levels in *B. tabaci*. Two TYLCV strains, TYLCV-Mld and TYLCV-IL, were transmitted to the same tomato cultivar using a single whitefly cohort (Fig. 9A). Thirty days post-inoculation, both strains induced typical disease symptoms, with TYLCV-IL causing significantly more severe symptoms than TYLCV-Mld. Viral titers in the intestines of viruliferous whiteflies were significantly (*F* = 168.16; df = 1, 32; *P* < 0.0001) higher in TYLCV-IL-infected individuals compared to TYLCV-Mld, as determined by immunofluorescence and qPCR analyses (Fig. 9B). This difference was evident as early as 2 days post-infection, coinciding with significantly higher *Bt-CYP6* expression in TYLCV-IL-infected whiteflies (Fig. 9C). In contrast, expression of *Bt-sEH2* and *Bt-sEH3* was more strongly suppressed in TYLCV-IL-infected whiteflies than in those infected with TYLCV-Mld.

**Fig 9.**
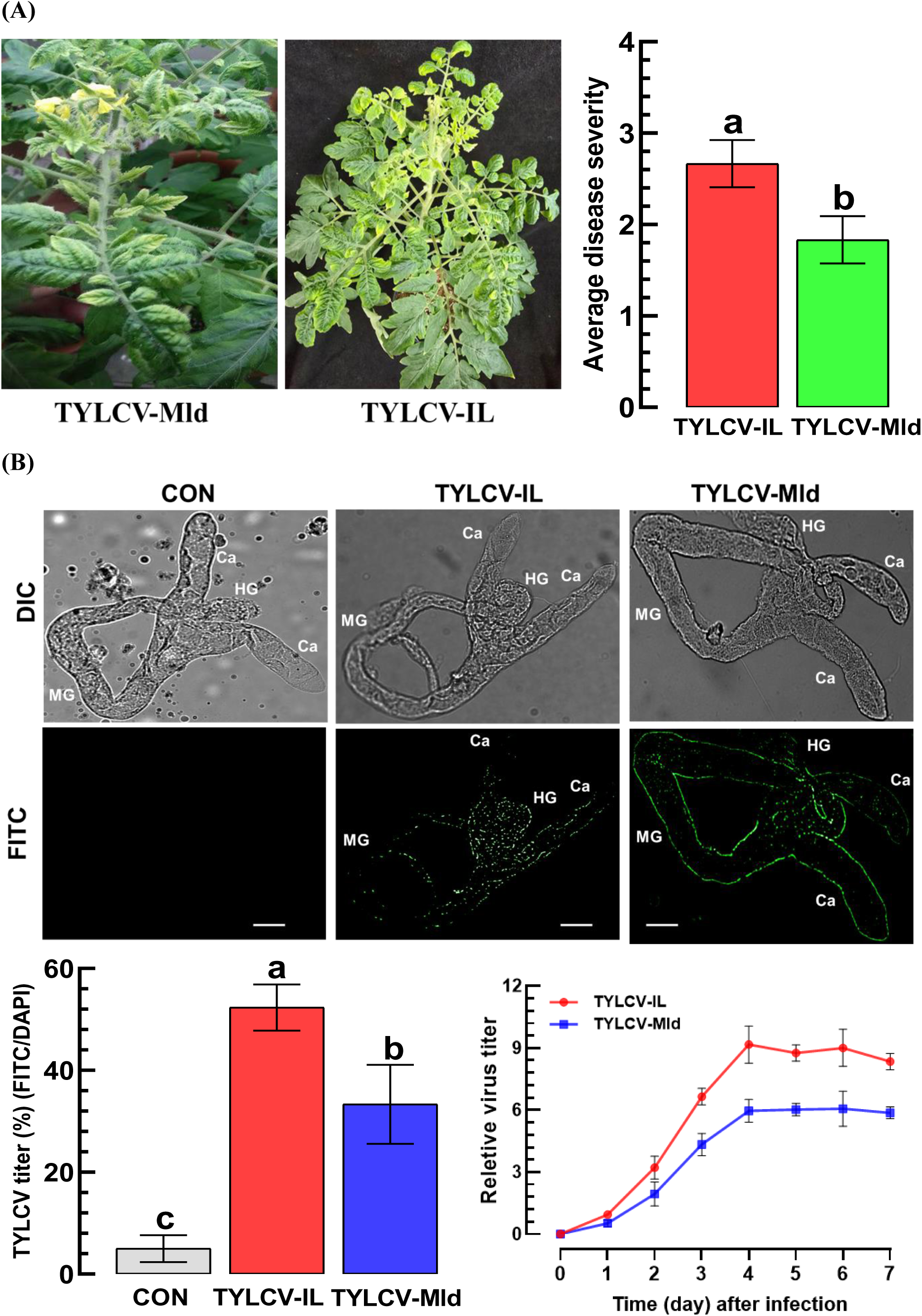
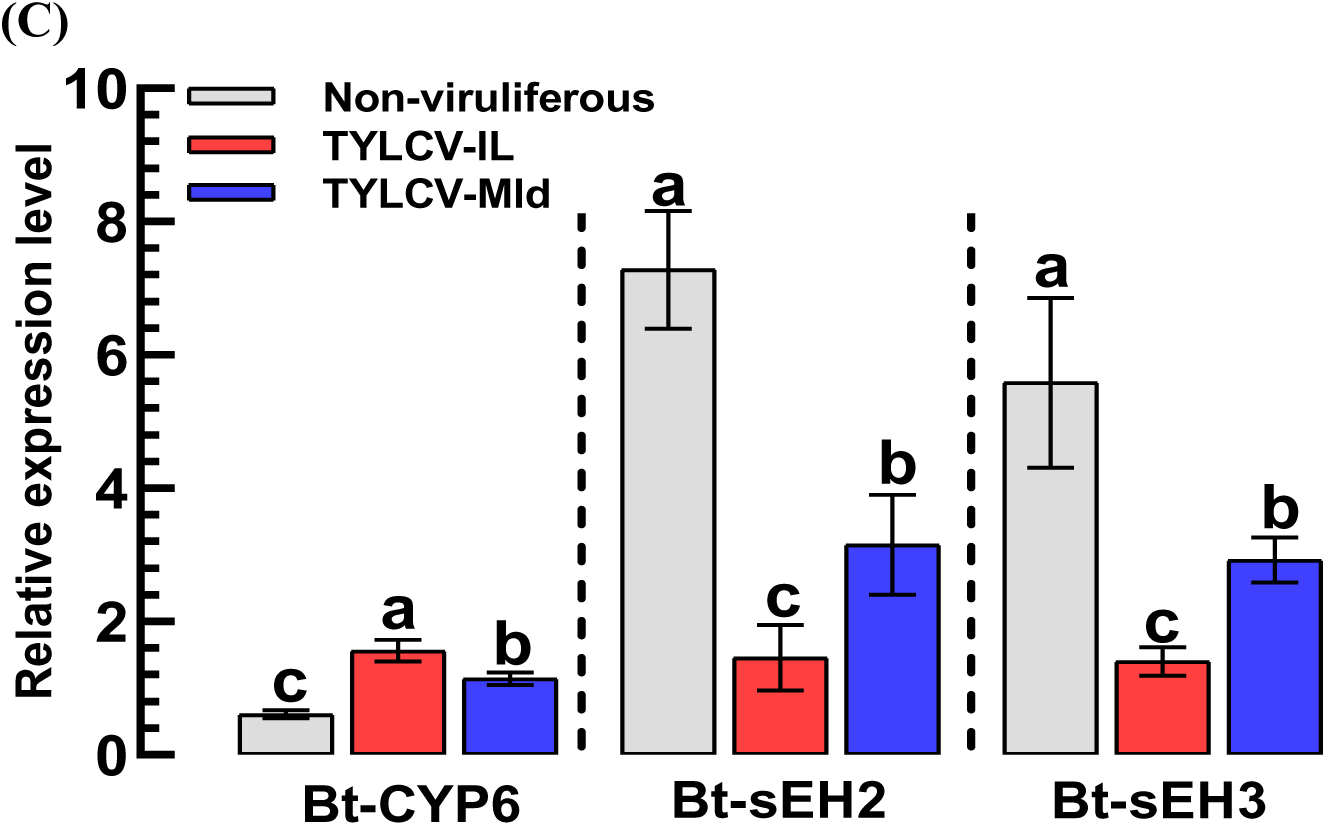
Differential control of EpOME synthetic machinery expression by two TYLCV strains (TYLCV-IL and TYLCV-Mld). (A) Difference in virulence between the two viral strains with respect to disease symptoms and severity after 30 days post-infection. Viruliferous *B. tabaci* after 48 h AAP were used to inoculate healthy tomato plants. (B) TYLCV multiplication in the intestine of *B. tabaci* by FISH analysis under differential interference contrast (‘DIC’) and fluorescein isocyanate (‘FITC’): ‘Ca’ for caeca, ‘MG’ for midgut, and ‘HG’ for hindgut. Scale bars represent 0.1 mm. Viral titers were assessed in FITC intensity normalized by the DAPI signal intensity in the intestine. In addition, the viral titers were assessed by qPCR from the whole-body samples for seven days post-infection. Each treatment was replicated three times. (C) Differential effect of the two TYLCV strains on modulating expressions of EpOME synthetic (*Bt-CYP6*) and degrading (*Bt-sEH2* and *Bt-sEH3*) genes. Different letters above standard deviation bars indicate significant differences among means at Type I error = 0.05 (LSD test).

### Apoptosis in the intestine of *B. tabaci* as a proviral response to TYLCV infection

TYLCV infection induced apoptosis in the intestine, particularly in the midgut, as detected by TUNEL assays (Fig. 10A). Apoptotic signals were reduced by PGE_2_ supplementation but enhanced by 12,13-EpOME treatment in viruliferous whiteflies. Expression levels of two apoptosis-associated caspases, the initiator caspase *Dronc* and the effector caspase-1 (*Cas1*), were significantly upregulated following TYLCV infection (Fig. 10B), with higher induction observed in TYLCV-IL-infected whiteflies compared to TYLCV-Mld. Functional involvement of apoptosis in viral accumulation was further supported by treatment with the apoptosis inhibitor Z-VAD, which significantly reduced viral titers in whiteflies infected with either strain (Fig. 10C).

**Fig 10.**
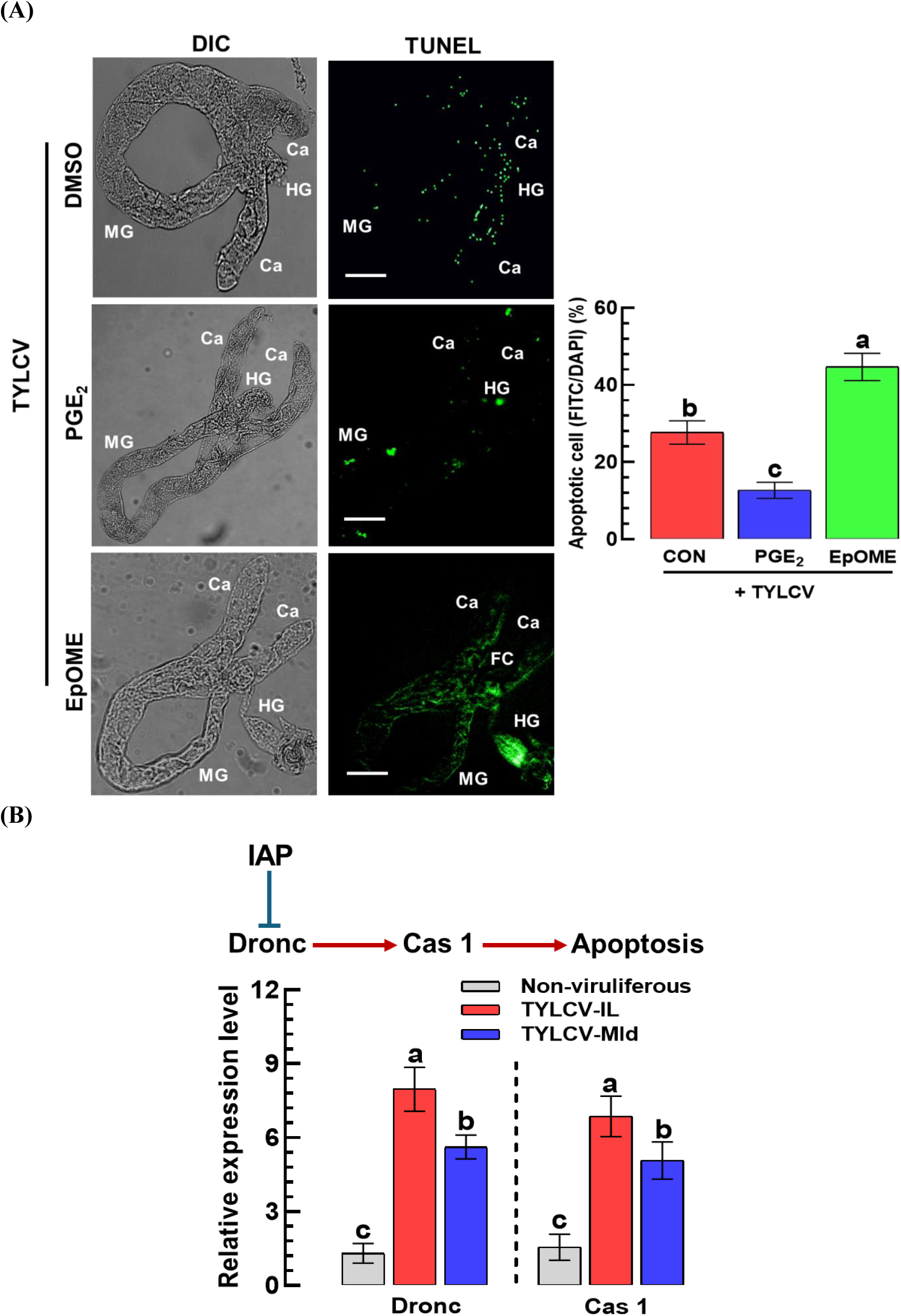
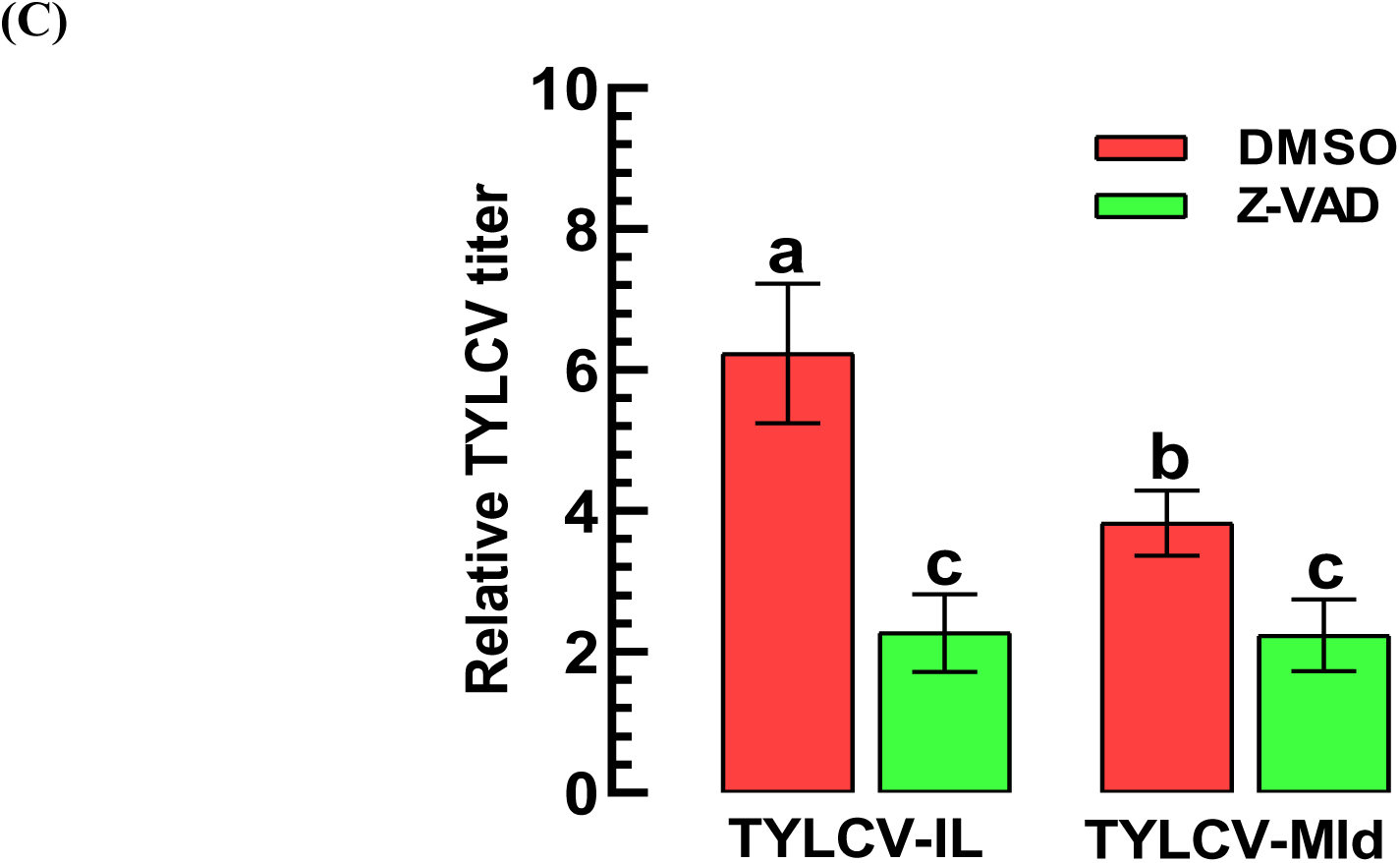
Apoptosis in the intestine of *B. tabaci* as a proviral response to TYLCV. (A) Opposite roles of two oxylipins (PGE_2_ and EpOME, each 0.1 µg/mL) in inducing intestinal apoptosis of *B. tabaci* upon TYLCV-IL infection assessed by a TUNEL assay. The intestine was visualized at 24 h after viral infection using differential interference contrast (‘DIC’), and apoptotic signals were detected using fluorescein isothiocyanate (‘FITC’) under a fluorescence microscope (DM2500; Leica, Wetzlar, Germany): ‘Ca’ for caeca, ‘MG’ for midgut, and ‘HG’ for hindgut. Scale bar = 0.1 mm. Viral titers were assessed as FITC intensity normalized by the intensity of DAPI signal intensity in the intestine. ‘DMSO’ represents a solvent control. (B) Differential effect of two TYLCV strains on the expressions of an initiator (‘Dronc’, a target of inhibitor of apoptosis (‘IAP’)) and an effector (‘Cas 1’) caspases in viruliferous *B. tabaci*. (C) Influence of an apoptosis inhibitor (‘Z-VAD’, 50 μM) on the viral titer of TYLCV-IL or TYLCV-Mld in viruliferous *B. tabaci*. Relative TYLCV titers were measured by qPCR at 24 h post-infection. Different letters above standard deviation bars indicate significant differences among means at Type I error = 0.05 (LSD test).

### Immune-associated genes of *B. tabaci* function as either proviral or antiviral factors during TYLCV infection

Six immune-associated genes that were upregulated following TYLCV infection were selected for functional analysis (Fig. 11). EpOME treatment induced the expressions of *defensin*, *cathepsin-B*, and *cathepsin-L*, while it suppressed the expressions of *knottin*, *PGRP*, and *cathepsin-F* (Fig. 11A). Conversely, PGE_2_ treatment showed expression patterns opposite to those induced by EpOME, consistent with their antagonistic roles in regulating TYLCV infection. To clarify this expression control, individual RNAi treatments specific to each of the immune genes were performed and evaluated for the control of the viral titers (Fig. 11B). RNAi-mediated suppression of *defensin*, *cathepsin-B*, or *cathepsin-L* significantly reduced viral titers in viruliferous whiteflies, indicating proviral roles. In contrast, knockdown treatments of *knottin*, *PGRP*, or *cathepsin F* significantly increased viral titers, suggesting antiviral functions.

**Fig 11.**
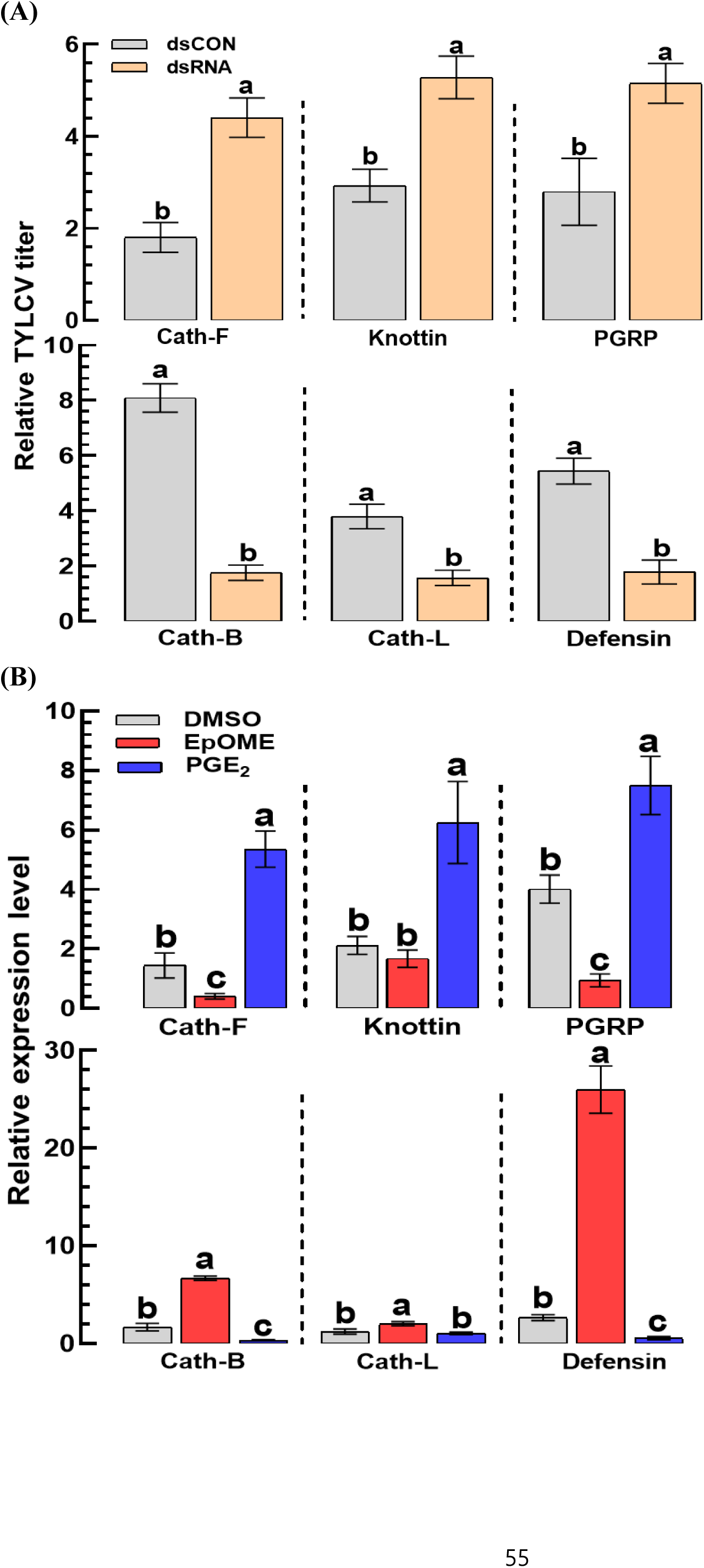
Selective control of proviral and antiviral immunity-associated genes of *B. tabaci* by EpOME. (A) Influence of individual RNAi treatments specific to six immunity-associated genes (*Defensin*, *knottin*, *PGRP*, *cathepsin B* (*Cath-B*), *cathepsin F* (*Cath-F*), and *cathepsin L* (*Cath-L*)) on the TYLCV titer in viruliferous *B. tabaci*. Individual dsRNA (500 µg/mL) against each of the immune-associated genes of *B. tabaci* were administered to adults by feeding along with TYLCV-IL. At 24 h post-infection, TYLCV titer was measured by qPCR. (B) Influence of 12,13- EpOME (0.1 µg/mL) or PGE_2_ (10^-7^ M) addition on the expressions of the six immune-associated genes at 24 h post-treatment in viruliferous *B. tabaci*. Different letters above standard deviation bars indicate significant differences among means at Type I error = 0.05 (LSD test).

### A viral factor, C2, modulates host vector in up-regulating EpOME level upon TYLCV infection

To identify the virulence factor derived from TYLCV responsible for the up-regulation of host EpOME levels, six viral genes (V1, V2, C1, C2, C3, and C4; Fig. 12A) encoded in the TYLCV genome were functionally evaluated using an RNAi-based loss-of-function approach. Adult whiteflies infected with TYLCV were individually treated with gene-specific dsRNAs (dsRNA^V1^, dsRNA^V2^, dsRNA^C1^, dsRNA^C2^, dsRNA^C3^, and dsRNA^C4^). All RNAi treatments efficiently suppressed their corresponding target genes for at least 48 h post-infection (Fig. S2). Under these knockdown conditions, the expression levels of the EpOME biosynthetic gene (*Bt-CYP6*) and the EpOME degradation genes (*Bt-sEH2* and *Bt-sEH3*) were quantified (Fig. 12B). Among all viral genes tested, silencing of *TYLCV-C2* alone disrupted EpOME pathway regulation. Specifically, RNAi specific to *TYLCV-C2* markedly suppressed the TYLCV-induced up-regulation of *Bt-CYP6*, whereas knockdown of other viral genes had no significant effect. In addition, *TYLCV-C2* silencing prevented the virus-associated down-regulation of *Bt-sEH2* and *Bt-sEH3* expression, in contrast to the other RNAi treatments. These results indicate that TYLCV-C2 is the principal viral factor responsible for promoting EpOME accumulation by simultaneously enhancing EpOME biosynthesis and suppressing EpOME degradation in TYLCV-infected whiteflies. To further validate the role of TYLCV-C2-mediated EpOME signaling in viral infection, a rescue experiment was performed by orally supplying 12,13-EpOME to the TYLCV-C2-silenced viruliferous whiteflies. Exogenous 12,13-EpOME effectively restored viral accumulation to levels comparable to the dsCON control (Fig. 12C), confirming the role of TYLCV-C2 in host-mediated EpOME up-regulation to facilitate viral infection.

**Fig 12.**
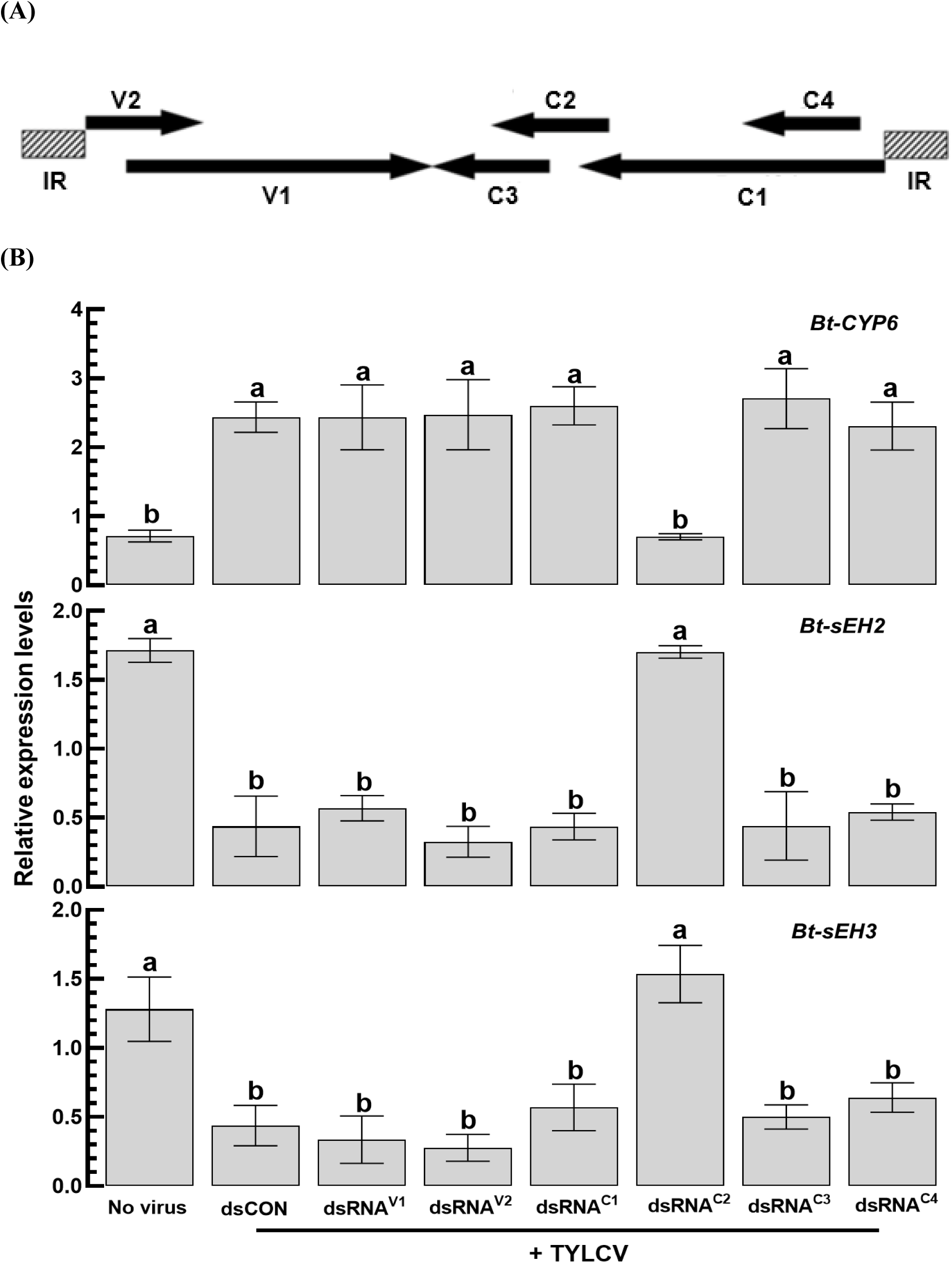
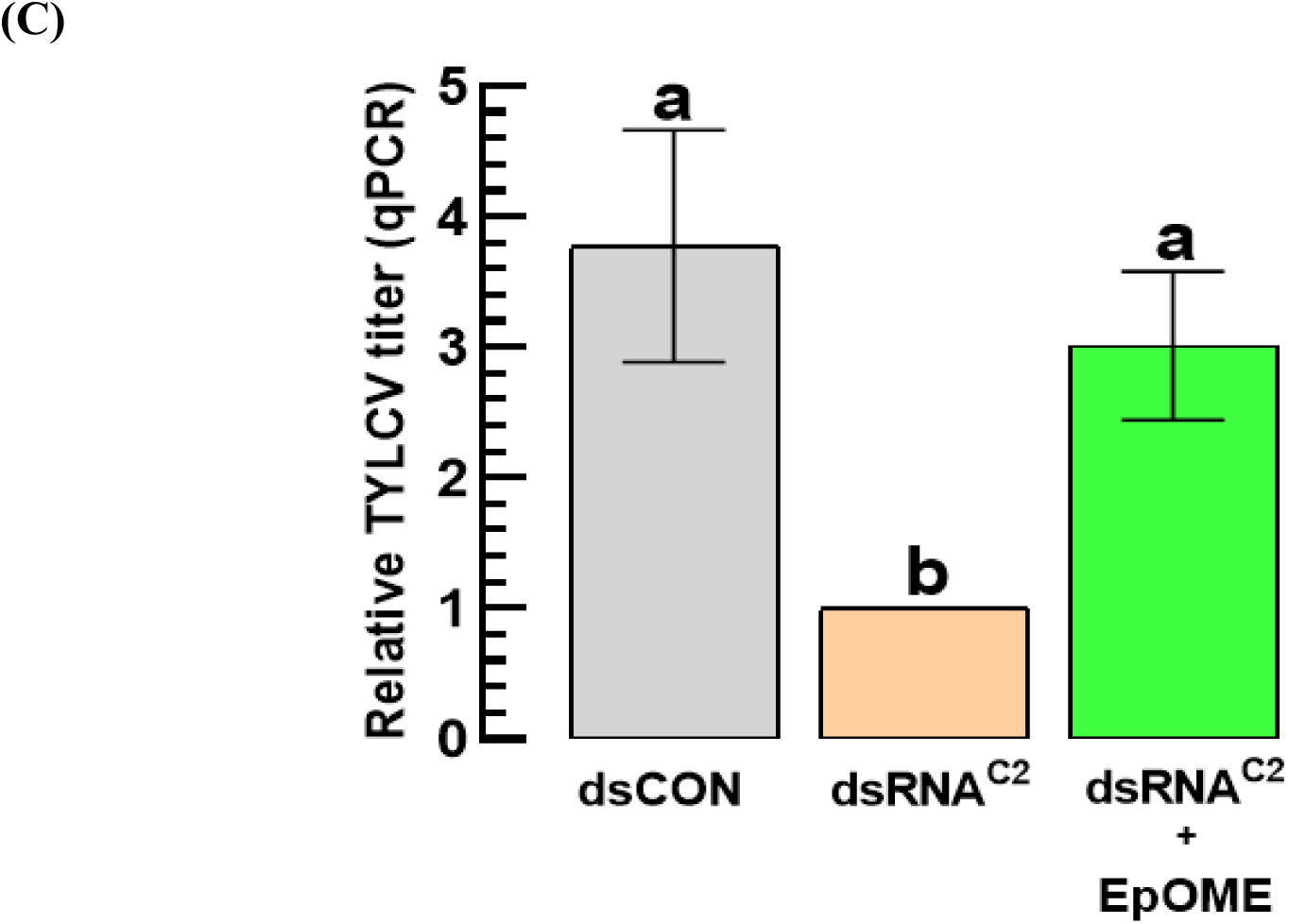
TYLCV-C2 as a virulent factor for manipulating host EpOME biosynthesis in *B. tabaci*. (A) TYLCV genome indicating six viral genes between intergenic region (IR) at both ends. (B) Effects of individual RNAi treatments (dsRNA^V1^, dsRNA^V2^, dsRNA^C1^, dsRNA^C2^, dsRNA^C3^, and dsRNA^C4^) specific to six viral genes, on the expression levels of *Bt-CYP6*, *Bt-sEH2*, and *Bt-sEH3* at 24 h post-RNAi treatment. ‘dsCON’ indicates a control dsRNA. TYLCV-IL was administered to adults, and the gene expression levels were assessed by qPCR. (C) Rescue effect of the suppressed viral titer induced by RNAi specific to TYLCV-C2 by EpOME addition. Each treatment was replicated three times. Distinct letters above the standard error bars signify significant differences among the means at a Type I error rate of 0.05 (LSD test).

## DISCUSSION

It has been recognized that TYLCV is not a passive passenger within its vector but actively reshapes whitefly development, immunity, and behavior to optimize its own transmission (27). TYLCV infection alters whitefly life-history traits, feeding behavior, and stress tolerance, illustrating a multifaceted strategy of vector manipulation. Despite these advances, the molecular mechanisms linking viral sensing in the gut to a durable, system-wide permissive state within the vector have remained poorly defined. Here, we demonstrate that TYLCV achieves this permissive state by rewiring vector oxylipin signaling, shifting the balance from antiviral eicosanoids toward proviral EpOMEs. In addition, this study identifies the viral factor responsible for this endocrine signal change.

This study demonstrated a progressive infection of TYLCV in the host intestine. Initially, the infection was observed in the midgut and hindgut, but not in the foregut. Later, the infection foci increased in number. Furthermore, the viral titers increased in a whole-body sample as well as in the intestine, supporting active viral replication of TYLCV in the vector insect. Although TYLCV was previously classified as a circulative, non-propagative virus transmitted by the whitefly *B. tabaci*, accumulating evidence indicates that TYLCV can replicate within its vector in a circulative–propagative transmission mode (28, 29). During this viral multiplication, TYLCV must have overcome defensive barriers derived from the whiteflies. Successful circulative transmission of TYLCV requires that the virus traverse multiple anatomical and physiological barriers while preserving vector viability (27, 30). At least three major anatomical barriers of the whiteflies restrict TYLCV movement: the midgut epithelial cells for entry and exit, the hemocoel, and the salivary gland cells for entry and exit (31). Viral entry into the midgut epithelium is mediated by the CUBAM receptor complex, composed of cubilin and amnionless, and proceeds through clathrin-dependent endocytosis (32). Following receptor binding, TYLCV is internalized into endosomal compartments, where low pH promotes virion dissociation from the receptor. Subsequent trafficking decisions determine whether virions are degraded in lysosomal compartments or transported across the basal membrane, a process involving sorting nexins such as Snx12 (31, 32). The resulting free virions in the midgut epithelial cells may transform host cells into favorable conditions for viral replication.

One of the physiological changes after viral infection was apoptosis in the midgut epithelium of the viruliferous *B. tabaci*. This phenomenon has also been reported in previous studies showing that TYLCV activates apoptotic pathways in *B. tabaci* and that experimental manipulation of apoptosis can alter viral accumulation, supporting a functional link between apoptosis and transmission biology (33, 34). Our findings extend this framework by demonstrating that intestinal apoptosis intensity is positively correlated with viral titer. This positive relation between apoptosis and TYLCV titer was further supported by treatment with the apoptosis inhibitor, Z-VAD, which led to a reduction in the viral titer. Furthermore, this study demonstrated that apoptosis in the gut is enhanced by EpOME signaling while being suppressed by eicosanoid signaling. The effect of EpOME treatment on gut apoptosis was further supported by the induction of caspase genes associated with the initiation and execution of apoptosis. Although apoptosis is classically antiviral in many insect–virus systems, including other plant viruses such as tomato spotted wilt virus (24), controlled epithelial turnover or barrier remodeling may facilitate virion movement across the gut wall or reduce intracellular degradation routes that would otherwise destroy virions. This interpretation is consistent with models in which endo-lysosomal processing represents a critical checkpoint determining viral degradation versus successful transcytosis (31). Recent studies of begomovirus–whitefly interactions emphasize that immune pathways such as apoptosis, autophagy, Toll, and JAK/STAT function as checkpoints that viruses must negotiate to establish persistent infections and maintain transmissibility (35). In parallel, detailed mechanistic studies have elucidated how TYLCV crosses the midgut barrier through receptor-mediated endocytosis and intracellular trafficking routes that balance transcytosis against degradation (31, 32). While these studies define where and how TYLCV moves within the vector, they do not fully explain how the internal immune-metabolic environment becomes permissive over time. Our results add a complementary layer by showing that TYLCV actively reprograms lipid mediator signaling, thereby influencing multiple immune and physiological checkpoints simultaneously. Specifically, we demonstrate that eicosanoid signaling suppresses TYLCV accumulation, whereas EpOMEs enhance viral titers. Indeed, TYLCV infection transcriptionally favors EpOME accumulation by inducing the EpOME synthase *Bt-CYP6* while suppressing EpOME-degrading sEH (*Bt-sEH2* and *Bt-sEH3*). Moreover, exogenous EpOME supplementation restores viral accumulation. Together, these results position oxylipin remodeling as an upstream coordinator of TYLCV-mediated vector manipulation. It was notable that remarkable changes in fatty acid metabolism were observed in DEG analysis between nonviruliferous and viruliferous whiteflies. This supports enhanced oxylipin production in response to TYLCV infection. EpOMEs and their corresponding DiHOMEs are well characterized in mammals as cytochrome P450–derived linoleate epoxides with immunomodulatory properties (36). In insects, EpOME biology has only recently begun to emerge. EpOMEs have been reported as immune suppressors in lepidopteran insects, and inhibition of soluble epoxide hydrolase can stabilize epoxy-fatty acids and mimic immunosuppressive phenotypes (17). Insect epoxide hydrolases are evolutionary diverse enzymes with functions extending beyond juvenile hormone metabolism to regulation of endogenous lipid mediators (37). Our study advances this literature by demonstrating EpOME-mediated immune regulation in a hemipteran vector and by directly linking EpOME accumulation to a plant virus transmission phenotype.

EpOME signaling acts upstream of immune effectors, selectively reshaping immune architecture rather than globally suppressing immunity. PGRPs function as upstream immune sensors and act as pattern-recognition receptors in insects and are implicated in antiviral defense signaling (38, 39). In our study, PGRP acted as an antiviral factor and was suppressed by EpOME signaling, suggesting that TYLCV dampens early immune sensing without disabling immune competence. Cathepsins displayed similar functional specialization: cathepsin-B and -L acted as proviral factors and were selectively induced by EpOME, whereas cathepsin-F exhibited antiviral activity and was suppressed, consistent with previous reports identifying cathepsin-B as a proviral determinant in *B. tabaci* (40, 41). Under EpOME-dominant conditions, *defensin* expression was enhanced and functioned as a proviral factor, whereas the cysteine-rich knottin peptide retained a clear antiviral role and was suppressed by EpOME but promoted by eicosanoid signaling. Importantly, knottin-1 has been previously shown to function as a critical antiviral restriction factor in *B. tabaci*, limiting TYLCV acquisition and transmission (42). Functional divergence among AMP classes has been reported in other insect–pathogen systems, where distinct AMPs contribute differently to resistance versus tolerance strategies (15, 43). Our findings extend this concept to TYLCV transmission.

Among viral genes, *TYLCV-C2* was identified as a viral determinant responsible for up-regulating EpOME signaling in the viruliferous *B. tabaci*. TYLCV-C2 promotes EpOME accumulation by inducing *Bt-CYP6* and suppressing *Bt-sEH2/3*, and the reduced viral accumulation in the TYLCV-C2 knockdown background could be rescued by exogenous EpOME application. TYLCV-C2 is a multifunctional virulence protein that subverts host ubiquitination pathway, suppress plant defenses and manipulate hormone-associated signaling, including jasmonic acid and salicylic acid pathways, to favor viral infection and vector performance (44, 45). A recent study further demonstrated that TYLCV-C2 alters salicylic acid–linked defense signaling in plants in ways that facilitate whitefly transmission (46). Our findings broaden the viral factor framework by showing that, in addition to its roles in the plant host, TYLCV-C2 directly reshapes the vector’s internal immune-metabolic environment.

Differences among TYLCV strains are often discussed in terms of symptom severity, viral accumulation, and epidemiological spread (5). However, mechanistic explanations for strain-specific differences in vector competence have largely focused on capsid–receptor interactions or intracellular trafficking efficiency (31, 32). Comparison of TYLCV-IL and TYLCV-Mld in this study reveals an additional mechanism: the more virulent TYLCV-IL strain induces stronger EpOME pathway activation, higher intestinal viral titers, and more pronounced apoptotic responses. This suggests that strain-specific capacity to reprogram oxylipin signaling contributes to vector competence, a mechanism that can operate in parallel with entry and trafficking pathways. Notably, our previous work demonstrated that wild-type and resistance-breaking strains of TSWV differentially upregulate EpOME biosynthesis in their thrips vector *F. occidentalis*, supporting a broader role for EpOME signaling in vector–virus interactions (47).

A survival–transmission balance has been proposed in which the JAK/STAT pathway protects whiteflies from TYLCV-associated fitness costs while permitting persistent infection, highlighting that vector immunity is not simply “on” or “off” but is optimized for coexistence with the virus (48). Our data aligns with this concept and extends it by identifying lipid mediators as rapid, systemic regulators linking metabolic status to immune outcomes. By increasing EpOME levels through C2 activity, TYLCV may shift the vector from an antiviral, eicosanoid-biased state toward a tolerant, EpOME-biased state that supports higher viral loads while preserving vector viability.

In conclusion, this study advances TYLCV transmission biology by identifying EpOMEs as endocrine-like mediators of vector competence, providing functional genetic evidence for specific EpOME biosynthesis and degradation genes, and positioning TYLCV-C2 as a direct regulator of vector lipid immunity. We propose a mechanistic model (Fig. 13) in which TYLCV-C2 shifts oxylipin signaling from an antiviral, eicosanoid-dominated state toward a proviral EpOME-dominated state, thereby coordinating apoptosis and immune effector specialization to create a system-wide permissive environment for persistent infection and transmission.

**Fig 13.**
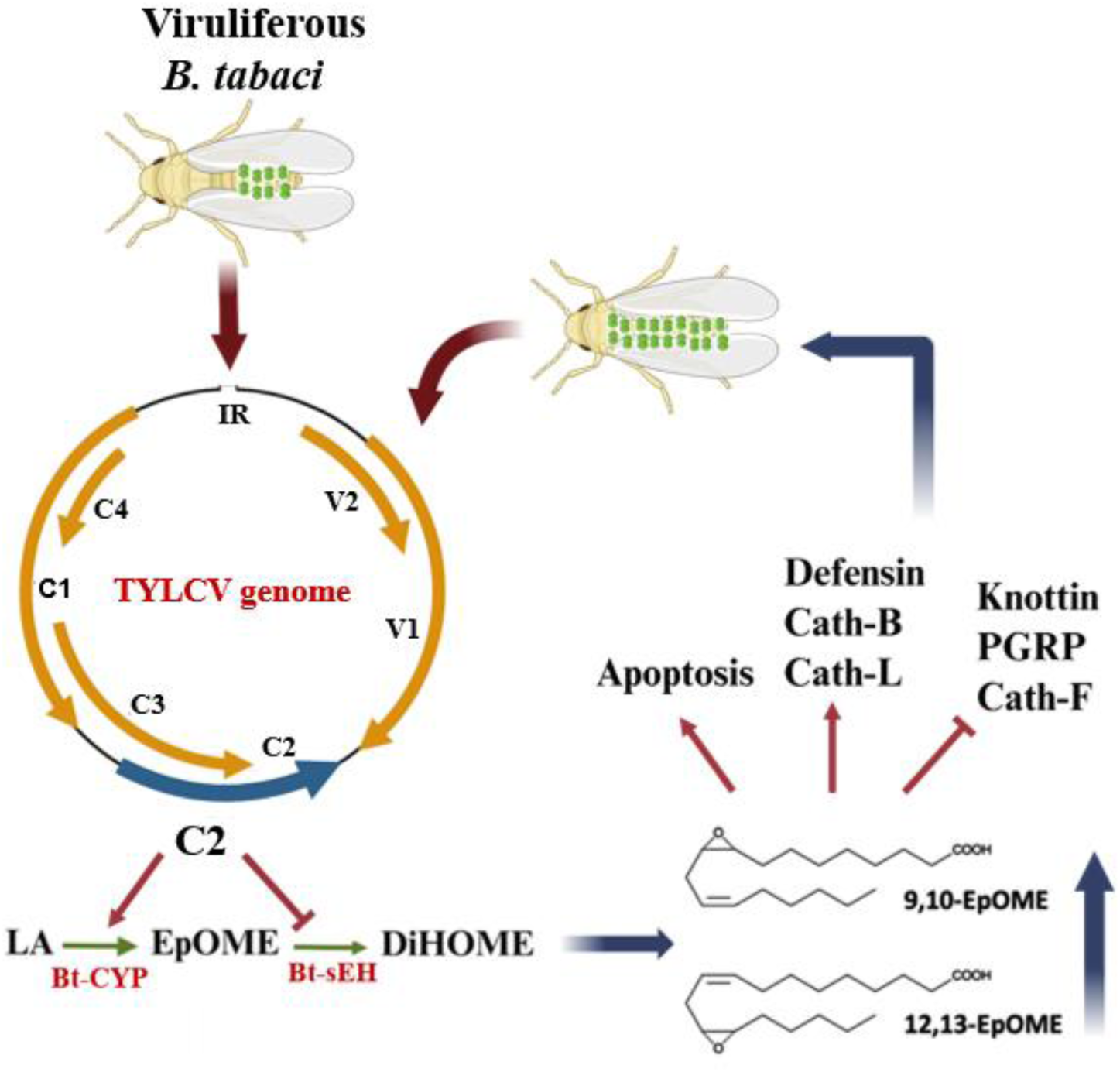
A working model of exploiting host EpOME for TYLCV multiplication in *B. tabaci*. Upon acquisition of TYLCV by the whitefly, TYLCV-C2 induces EpOME biosynthesis by inducing *Bt-CYP* expression while suppressing *Bt-sEH* expression. Up-regulated EpOME levels induce proviral factors but suppress antiviral factors, leading to a reshaping the physiological conditions of *B. tabaci* to facilitate TYLCV multiplication.

## MATERIALS and METHODS

### Rearing B. tabaci

A colony of the MED-type whitefly, *B. tabaci*, was originally collected from a greenhouse in Andong, Korea, and was maintained on tomato (*Solanum lycopersicum*) seedlings under controlled laboratory conditions (25 ± 1°C, 14 h light photoperiod and 70 ± 5% relative humidity). The whitefly strain was confirmed by sequencing a cytochrome oxidase I (COI) segment of the whitefly, which was deposited in GenBank under an accession number of PV993946.

### Origin of two TYLCV strains and disease severity index

To obtain virus-infected plants, infectious clones of TYLCV-IL (GenBank accession number: JN680149) or TYLCV-Mld (GenBank accession number: PX375450) strains were cloned into pCAMBIA1301 and inoculated into tomato seedlings by agro-inoculation. All plants were grown in insect-proof greenhouses under natural sunlight supplemented with artificial lighting for 14 h a day and a controlled temperature of 25 ± 3°C. The inoculated plants that showed typical symptoms of virus infection at 25 ∼ 30 days post inoculation were used for whitefly transmission. Virus infection of all inoculated plants was confirmed by PCR using specific primers (Table S1). The disease severity caused by the inoculated TYLCV strains on tomato plants was evaluated visually for 25 days post inoculation. Symptom development was scored using a disease index ranging from 0 to 4, where 0 = no symptoms and 4 = very severe symptoms (severe stunting, leaf curling, cupping, and growth arrest), following the method of Friedmann et al. (49). Intermediate scores (0.5, 1.5, 2.5, and 3.5) were used to improve the precision of disease severity assessment.

### TYLCV infection of *B. tabaci* and viral titer analysis

Newly emerged adult whiteflies (2 ∼ 4 days post-emergence) were collected in a batch of 50 individuals for an experimental unit and transferred onto tomato leaves infected with either the TYLCV-IL or TYLCV-Mld strain, placed inside large Petri dishes (100 × 40 mm, SPL Life Sciences, Seoul, Korea). After a 24-hour acquisition access period (AAP), whiteflies were collected and subjected to RNA and DNA extraction. To determine viral titer dynamics in infected *B. tabaci* over time, the TYLCV-infected whiteflies were released into small Petri dishes (50 × 15 mm, SPL Life Sciences), sealed with a double layer of plastic tape and provided with 30% sucrose for seven days. Each day, a group of 20 insects was collected for DNA extraction and quantitative PCR (qPCR) using specific primers targeting the TYLCV coat protein gene (Table S1).

### Test compounds and preparation of commonly used chemicals

9,10- and 12,13-EpOMEs, and prostaglandin E₂ (PGE₂) were purchased from Cayman Chemical (Ann Arbor, MI, USA). 12-(3-Adamantan-1-ylureido)dodecanoic acid (AUDA), benzyloxycarbonyl-Val-Ala-Asp-fluoromethyl ketone (Z-VAD), arachidonic acid (AA; 5,8,11,14-eicosatetraenoic acid), dexamethasone (DEX; (11β,16α)-9-fluoro-11,17,21-trihydroxy-16-methylpregna-1,4-diene-3-one), esculetin (ESC; 6,7-dihydroxycoumarin), and naproxen (NAP; (S)-(+)-2-(6-methoxy-2-naphthyl) propionic acid) were obtained from Sigma-Aldrich Korea (Seoul, Korea) and dissolved in dimethyl sulfoxide (DMSO) to prepare stock solutions. 5-Bromo-2′-deoxyuridine (BrdU) and anti-BrdU antibody were purchased from Abcam (Cambridge, UK). Terminal deoxynucleotidyl transferase, fluorescein isothiocyanate (FITC)-conjugated anti-mouse IgG antibody, and 4′,6-diamidino-2-phenylindole (DAPI) were obtained from Thermo Fisher Scientific (Wilmington, DE, USA). Bovine serum albumin (BSA), DMSO, and t-octylphenoxy-polyethoxyethanol (Triton X-100) were purchased from Sigma-Aldrich Korea. Phosphate-buffered saline (PBS) was prepared with 100 mM phosphoric acid containing 0.7% NaCl and adjusted to pH 7.4 with 1 M NaOH.

### RNA interference (RNAi) treatment to *B. tabaci* using double-stranded RNA (dsRNA)

Template DNAs were amplified by PCR using gene-specific primers (Table S1) containing the T7 promoter sequence at their 5′ ends, following previously described methods (26). The resulting PCR products were used as templates for *in vitro* transcription with the MEGAscript RNAi Kit (Ambion, Austin, TX, USA) according to the manufacturer’s instructions. The synthesized dsRNAs were combined with Metafectene PRO (Biontex, Munchen, Germany) at a 1:1 (v/v) ratio and incubated at room temperature (RT) for 30 min to form dsRNA–liposome complexes. Oral delivery of dsRNA was performed using a parafilm-based feeding apparatus prepared in sterile small Petri dishes. A synthetic diet containing 30% (w/v) sucrose and dsRNA at a final concentration of 500 ng/µL was applied to the inner surface of a stretched parafilm membrane. The membrane was lightly punctured at the feeding zone to enable stylet access. Adult *B. tabaci* (50 individuals per replicate) were placed on the outer side of the membrane and maintained under controlled conditions (25 ± 1°C, 70 ± 5% relative humidity, 14:10 h light:dark cycle) to allow ingestion of the dsRNA-containing diet. Gene silencing efficiency was evaluated by quantitative PCR within 48 h following exposure.

### Test compound treatment to *B. tabaci*

To treat the test compounds including AA (1 μg/mL), 12,13-EpOME (0.1 μg/mL), Z-VAD-FMK (50 μM), AUDA (10 μg/mL), DEX (1 μg/mL), NAP (1 μg/mL), ESC (1 μg/mL), PGE_2_ (0.1 μg/mL), PGD_2_ (0.1 μg/mL), and PGF_2α_ (0.1 μg/mL) to the adult whiteflies, 1 g of TYLCV-infected tomato leaves were homogenized with 10 mL of filter (0.22 μm pore size)-sterilized PBS (1:10, w/v) and centrifuged at 14,000× *g* for 5 min. The supernatant was used as the virus suspension and mixed with 30% (w/v) sucrose solution at a 1:1 ratio. Next, each compound was mixed with this mixture (to get the final concentration as mentioned before) and placed on the double layers of parafilm that covered the opening of the small Petri dish as explained above. After 24 h of feeding, groups of 50 whiteflies were collected for DNA extraction for virus titer analysis or RNA extraction for mRNA level analysis.

### Nucleic acid extraction, qPCR, and RT-qPCR

For transcript analysis, total RNA was isolated from the treated whiteflies using TRIzol reagent (Ambion) according to the manufacturer’s instructions. The concentration and purity of RNA samples were assessed using a NanoDrop spectrophotometer (Thermo Fisher Scientific). Reverse transcription was performed with 100 ng of total RNA using the RT-Premix kit (Intron Biotechnology, Seoul, Korea) containing an oligo(dT) primer, following the manufacturer’s protocol.

To quantify the viral titer, the treated whiteflies were homogenized in DNA extraction buffer (Biosearch Technologies, Seoul, Korea). The homogenate was incubated at 100°C for 5 min to lyse the cells, followed by centrifugation at 14,000× *g* for 3 min. The resulting supernatant was transferred to new tubes and used as the DNA template for subsequent experiments.

Gene expression levels and viral titers were determined by RT-qPCR and qPCR, respectively, using a Step One Plus Real-Time PCR System (Applied Biosystems, Singapore) and Power SYBR Green PCR Master Mix (Toyobo, Osaka, Japan) according to the manufacturer’s instructions. Each RT-qPCR reaction (20 µL total volume) contained 10 µL of 2× Power SYBR Green PCR Mix, 100 ng of DNA or cDNA template, and 10 pmol of each specific primer (Table S1). The *Bt-EFα* (Table S1) gene was used as an internal reference for normalization. Amplification specificity was confirmed by melting curve analysis. Each treatment was performed in triplicate using independent biological samples. Relative gene expression levels were calculated using the comparative Ct (2⁻^ΔΔCt^) method (50).

### Insect sample preparation to quantify EpOMEs

To determine EpOME levels in *B. tabaci* following TYLCV infection, lipids were extracted from both non-viruliferous and viruliferous adult whiteflies. After a 24-hour AAP on TYLCV-infected tomato plants, 200 adult whiteflies were collected and washed three times with chilled PBS. Each sample was subjected to three rounds of ultrasonic homogenization (1 min each) in PBS using an ultrasonicator (Bandelin Sonoplus, Berlin, Germany) set to 80% power. The pH of the homogenate was adjusted to 4.0 with 1 N HCl. Lipids were extracted by adding 1 mL of ethyl acetate, and the upper organic phase was separated. The remaining aqueous phase was re-extracted twice with ethyl acetate. Combined extracts were evaporated under a gentle nitrogen stream to a final volume of approximately 500 µL and loaded onto a small silicic acid column (2 × 90 mm; containing 30 mg of Type 60A silicic acid, 100–200 mesh; Sigma-Aldrich Korea). Sequential elusion was performed with solvents of increasing polarity in the order of 100% ethyl acetate, ethyl acetate/acetonitrile (1:1, v/v), acetonitrile/methanol (1:1, v/v), and finally 100% methanol. The ethyl acetate fraction was collected and used for EpOME quantification. Each treatment was performed in triplicate using independently prepared samples.

### LC-MS/MS analyses

EpOME quantification was performed using an LC–MS/MS system (QTrap 4500; AB Sciex, Framingham, MA, USA) equipped with an autosampler, binary pump, and column oven. Chromatographic separation was achieved on a C18 column (2.1 × 150 mm, 2.7 µm; Osaka Soda, Osaka, Japan) maintained at 40°C. The mobile phases consisted of 0.1% formic acid in water (solvent A) and 0.1% formic acid in acetonitrile (solvent B). The gradient program was as follows: 30% B (0–2 min), 30–65% B (2–12 min), 65–95% B (12–12.5 min), 95% B (12.5–25 min), and re-equilibration at 30% B (25–30 min). The flow rate was set to 0.4 mL min⁻¹, the injection volume was 10 µL, and the autosampler temperature was maintained at 5°C. Detection was performed using an electrospray ionization (ESI) source in negative ion mode. Optimized ion source parameters were as follows: temperature, 600°C; curtain gas, 32 L min⁻¹; ion gas, 60 L min⁻¹; and spray voltage, –4,000 V. Data were acquired in multiple reaction monitoring mode using nitrogen as the collision gas. Peak detection, integration, and quantification were conducted with MassView 1.1 software (AB Sciex).

### KEGG analysis of *B. tabaci* between viruliferous and nonviruliferous adults

RNA-seq data from non-viruliferous (BioSample: SAMN10060942; SRA: SRS3774498) and TYLCV-viruliferous *Bemisia tabaci* MED (BioSample: SAMN10060936; SRA: SRS3774502) were retrieved from the NCBI database. Raw reads were processed and analyzed using CLC Genomics Workbench (v24.0.1; QIAGEN, Hilden, Germany). Low-quality reads and adapter sequences were removed using the Trim Reads tool prior to mapping against the *B. tabaci* reference genome (GenBank accession no. GCF_918797505.1). Gene expression levels were quantified and normalized as transcripts per million (TPM) to enable comparisons between non-viruliferous and TYLCV-infected samples. Gene identifiers were assigned based on annotated gene symbols and corresponding LOC numbers obtained from the *B. tabaci* general feature format file. Differentially expressed genes (DEGs) between non-viruliferous and viruliferous groups were identified using the Differential Expression for RNA-Seq tool in CLC Genomics Workbench. Functional annotation was performed using Kyoto Encyclopedia of Genes and Genomes (KEGG) pathway analysis. Gene symbols were converted to KEGG gene IDs using the bioDBnet ID conversion tool, and pathway mapping was conducted with KofamKOALA (KEGG; https://www.genome.jp/tools/kofamkoala). KEGG pathway enrichment analysis was used to identify metabolic and functional pathways significantly affected by TYLCV infection based on differential TPM profiles. Data visualization was performed using R software (version 4.3.1) with the ggplot2 package.

### Bioinformatics to predict eicosanoid- and EpOME-associated genes from *B. tabaci* genome

To predict *PLA_2_*, *CYP*, and *sEH* genes of *B. tabaci*, all annotated amino acid sequences of these gene families were retrieved from the NCBI-GenBank database (www.ncbi.nlm.nih.gov) and aligned with their orthologs from other representative insects and vertebrate species (Table S2). Multiple sequence alignments were conducted using ClustalW within MEGA11 (51). Their phylogenetic trees were constructed using the Neighbor-joining method, with 1,000 replicates to calculate bootstrap values, employing MEGA11.

### Fluorescence in situ hybridization (FISH) analysis

After 24 h-feeding on the TYLCV-infected diet, adult whitefly midguts were dissected on sterilized glass slides and fixed with 4% paraformaldehyde for 1 h at RT. Following fixation, the tissues were rinsed with 1× PBS and permeabilized with 1% Triton X-100 in PBS for 1 h at RT. After an additional PBS wash, the samples were rinsed in 2× saline-sodium citrate (SSC) buffer and incubated at 42°C with 25 μL of pre-hybridization buffer containing 2 μL of yeast tRNA, 2 μL of 20× SSC, 4 μL of dextran sulfate, 2.5 μL of 10% SDS, and 14.5 μL of deionized H₂O. The incubation was carried out for 1 h under dark and humid conditions. The pre-hybridization buffer was then replaced with a hybridization buffer consisting of 5 μL of deionized formamide and 1 μL of fluorescein-labeled oligonucleotide probe in 19 μL of the initial pre-hybridization mix. DNA oligonucleotide probes targeting the TYLCV *cp* gene (Table S1) were labeled at the 5′ end with fluorescein amidite (FAM) and purified by high-performance liquid chromatography (Bioneer, Daejeon, Korea). Slides were covered with RNase-free coverslips and incubated overnight (18 h) in a humid chamber at 42°C. Following hybridization, the gut tissues were washed twice with 4× SSC for 10 min each, followed by a 5 min wash in 4× SSC containing 1% Triton X-100 at RT. The samples were then washed three times with 4× SSC, incubated at 37°C for 30 min in 1% anti-rabbit FITC-conjugated antibody (Thermo Fisher Scientific) in PBS, and protected from light. After two washes with 4× SSC (10 min each) and one wash with 3× SSC, the midgut samples were air-dried. A drop of 50% glycerol was added, and after a 15 min incubation at RT, the samples were mounted with a coverslip and examined under a fluorescence microscope (DM2500, Leica, Wetzlar, Germany) at 200× magnification.

### Terminal deoxynucleotidyl transferase dUTP nick end labeling (TUNEL) assay

TUNEL assays were performed using the *in-situ* Cell Death Detection kit (Abcam, Cambridge, UK). Adult whitefly midguts were dissected in PBS and placed on 22 × 22 mm coverslips. The tissues were then incubated with 10 μM BrdU and terminal transferase for 1.5 h. Following incubation, samples were fixed in 4% paraformaldehyde for 1 h at RT, rinsed with PBS, and permeabilized with 0.3% Triton X-100 in PBS for 2 h. After permeabilization, tissues were blocked for 1 h with 5% BSA in PBS and subsequently incubated with an anti-BrdU antibody (1:15 dilution in blocking buffer) for 1 h at RT. Unbound antibodies were removed by three PBS washes, followed by incubation with an FITC-conjugated anti-mouse IgG secondary antibody (1:300 dilution in blocking buffer) for 1 h. After three additional PBS washes, 10 μL of DAPI solution (1:1,000 dilution in blocking buffer) was added and incubated for 5 min at RT. Samples were rinsed with PBS, mounted with 10 μL of a glycerol–PBS mixture, and covered with a glass coverslip. Fluorescence signals were observed using a Leica DFC450C fluorescence microscope in FITC mode. Each experimental treatment was performed in triplicate.

### Statistical analysis

All the continuous variable data were analyzed by one-way analysis of variance (ANOVA) with the PROC GLM procedure in the SAS program (SAS Institute, 1989). Means were compared using the least significant difference (LSD) test at Type I error = 0.05. All the graphs in this study were prepared using GraphPad Prism v.8.0.1 (Boston, MA, USA).

## Acknowledgments

This work was supported by a grant (No. 2022R1A2B5B03001792) from the National Research Foundation (NRF) funded by the Ministry of Science, ICT and Future Planning, Republic of Korea. It was also supported by the research grant from a Glocal Project of Gyeongkuk National University.

